# Comprehensive characterization of *Plasmodium vivax* antigens using a high-density peptide array

**DOI:** 10.64898/2026.03.17.712326

**Authors:** Rosita R. Asawa, Brittany Hazzard, Kieran Tebben, John Tan, Tineke Cantaert, Andrea Berry, Niraj Tolia, Jean Popovici, David Serre

**Affiliations:** Institute for Genome Sciences, University of Maryland School of Medicine, Baltimore MD, USA; Nimble Therapeutics Inc., Madison WI, USA; Immunology Unit, Institut Pasteur du Cambodge, Phnom Penh, Cambodia; Center for Vaccine Development, University of Maryland School of Medicine, Baltimore MD, USA; Laboratory of Malaria Immunology and Vaccinology, National Institute of Allergy and Infectious Diseases, Bethesda MD, USA; Malaria Research Unit, Institut Pasteur du Cambodge, Phnom Penh, Cambodia; Infectious Disease Epidemiology and Analytics G5 Unit, Institut Pasteur, Paris, France

## Abstract

*Plasmodium vivax* is the second most prevalent *Plasmodium* species, with 2.5 billion people at risk of infection worldwide and around 10 million cases of clinical vivax malaria every year. Despite the clinical importance of this pathogen, very little is known about the *P. vivax* proteins recognized by the host immune system, which hinders our ability to select vaccine candidates or develop efficient serological markers. To comprehensively characterize immunogenic *P. vivax* proteins, we designed a high-density peptide array containing 4.2 million peptides covering the entire protein sequence of all *P. vivax* genes and analyzed antibody responses of infected and malaria-naïve individuals. We identified a total of 283 proteins that are commonly immunogenic in symptomatic individuals. These proteins included most proteins known to be involved in erythrocyte invasion, a putative new invasion protein, several nucleoporins, and many uncharacterized proteins that should be further investigated for their roles during blood-stage infections. These analyses also revealed a unique pattern of antibody response against PIR proteins in asymptomatic individuals, that could be associated with protection against clinical vivax malaria. Overall, these data provide an agnostic and comprehensive perspective on immunogenic *P. vivax* proteins and constitute an important resource for the malaria community to develop new tools for better detecting and eliminating this important human pathogen.

## Introduction

Malaria is an infectious disease that affects millions of individuals each year (1). It is caused by unicellular parasites of the *Plasmodium* genus, with *Plasmodium falciparum* and *Plasmodium vivax* responsible for more than 95% of the cases worldwide. *P. vivax* is the second most prevalent *Plasmodium* species, with 2.5 billion people at risk of infection and 10 million vivax malaria cases each year, primarily in Southeast Asia, India, and South America (1). While *P. vivax* and *P. falciparum* both cause human malaria, these parasites have very different biological features (2, 3). For example, vivax malaria typically has a lower parasitemia compared to falciparum malaria, in part due to the preferential invasion of immature reticulocytes by *P. vivax* parasites (4, 5) (while *P. falciparum* parasites can invade any red blood cell), and a lower pyrogenic threshold (6). In addition, *P. vivax* parasites have a hepatic stage not observed in *P. falciparum*: the hypnozoites. Hypnozoites can remain dormant in the liver for weeks to months after the initial infection has been cleared before reactivating to cause new blood stage infections (7). This dormant stage allows *P. vivax* parasites to persist in areas where mosquitoes are present only seasonally, broadening the geographical distribution of the parasites, and facilitates parasite circulation, which complicates elimination and eradication efforts (8).

The limited sensitivity and specificity of the rapid diagnostic tests currently available for *P. vivax* and the lack of biomarkers for identifying latent hypnozoite burdens have limited our ability to efficiently identify and treat *P. vivax* infections (9, 10). Additionally, the lack of an *in vitro* culture system for *P. vivax* dramatically limits the scope of laboratory investigations that can be conducted on this important human pathogen. As a consequence, the therapeutic toolkit available to treat and prevent *P. vivax* infections remains limited and is primarily derived from drugs developed against *P. falciparum*. Artemisinin-based combination therapy and, in some regions, chloroquine are the current standard-of-care for the treatment of vivax malaria (1), but these treatments only target blood-stage parasites, leaving hypnozoites unaffected and able to cause relapses after successful treatment of the blood-stage infection (9, 11). Only primaquine and tafenoquine effectively target hypnozoites but the use of these drugs in endemic areas is complicated by severe side effects in G6PD-deficient individuals, a common trait geographically correlated with historical exposure to malaria (12). Finally, while several promising vaccines against *P. falciparum* are in development (13), vaccine research on *P. vivax* has lagged behind, partially due to our limited understanding of *P. vivax* antigens: most studies thus far have focused on a handful of selected candidates primarily derived from the *P. falciparum* literature, with the exception of the Duffy binding protein (DBP) (14, 15). Unfortunately, the *P. vivax* vaccine candidates that are currently in early-stage clinical trials seem to only confer partial protection (16–21), highlighting the need to identify novel, evidence-based, candidates.

During a malaria infection, the host immune system mounts a response against the parasites, which involves the expansion of antigen-specific B and T cells as well as the production of antibodies against specific parasite proteins (22, 23). Upon repeated infections, this response leads to the acquisition of immunity against the disease, if not sterilizing (24). Understanding which parasite proteins are recognized by the immune system and elicit the production of a robust antibody response is key, not only to identifying new targets for vaccines, but also to developing biomarkers against specific stages (*e.g.*, the dormant hypnozoites) or of recent *Plasmodium* exposure (25–27). However, the identification of antigenic molecules is complicated by the fact that *Plasmodium* parasites express thousands of different proteins at different times during their complex life cycle (28, 29). In addition, high-throughput synthesis of recombinant proteins for testing in serological assays is complicated by the complex structure of eukaryotic genes, which often contain introns (and can sometimes be transcribed into different isoforms (30)). As a consequence, we know little about the antibody reactivity of most *P. vivax* proteins and most studies so far have focused on a handful of candidate proteins selected for their biological role (*e.g.*, in red blood invasion) (25, 31, 32). A few large screens have been conducted using protein arrays, but the difficulty and costs of synthesizing *Plasmodium* proteins have limited their scope to, at best, a few hundred genes that are expressed mainly in blood stage parasites (33–35). High-density peptide arrays are an exciting alternative for simultaneously characterizing antibody responses to thousands of antigens and, potentially, for identifying their specific epitopes (36). The synthesis of millions of short peptides (typically 10 - 20 amino acid long capturing linear B-cell epitopes (37, 38)) is now possible, enabling screening of the entire protein repertoire of pathogens. Such peptide array technology has notably been successfully applied to *P. falciparum* to i) identify linear immunogenic B-cell epitopes in individuals naturally exposed to malaria (39–41), ii) measure serological responses post-vaccination (42, 43), and iii) characterize the strain specificity of antibody responses (42, 44).

Here, we describe the design and analysis of a proteome-wide *P. vivax* high-density peptide array and characterize the antibody response of Cambodian individuals during a *P. vivax* infection. We identified 283 proteins that elicit antibody responses across participants, including proteins known for their role in red blood cell invasion as well as many proteins with unknown functions that could play important roles in host/parasite interactions.

## Materials and Methods

### Study population

Ten serum samples were collected in 2014 during a drug efficacy study in the Ratanakiri Province in Northeastern Cambodia (45). Febrile patients with a positive non-falciparum rapid diagnostic test were enrolled and, after confirmation by PCR that the infection was caused solely by *P. vivax*, serum was collected from each participant before antimalarial treatment (**Supplemental Table 1**). In addition, we analyzed sera from five Cambodian individuals, enrolled in a longitudinal study in the Mondolkiri Province (46), that were infected with *P. vivax* but were asymptomatic at the time of the collection (**Supplemental Table 1)**. As negative controls, we analyzed serum from ten North American individuals living in Baltimore (USA) and with no history of malaria exposure, no travel to or residence in a country where malaria is endemic, no history of sickle cell trait, no vaccines received in the past year, no current acute illness, and no history of fever in the three days prior to blood draw.

### Ethics statement

All Cambodian samples were obtained after written informed consent from the participants, and the analyses were approved by the National Ethic Committee of the Cambodian Ministry of Health (#038NECHR, 239NECHR and 057-NECHR). Serum collection from the North American individuals was approved by the University of Maryland, Baltimore Institutional Review Board (Protocol # HP-00071211).

### High-density peptide array design and data generation

We designed a high-density array containing 4.2 million peptides covering the entire amino acid sequence of all protein-coding genes annotated in the *P. vivax* P01 genome (47) (retrieved from PlasmoDB version 37). Briefly, we bioinformatically split the amino acid sequences from all annotated *P. vivax* protein-coding genes into 16 amino acid peptides overlapping the next peptide in the protein sequence by 15 amino acids (**Figure 1** and **Figure S1**). After removing redundant peptides (*i.e.*, we kept only one instance of a peptide found in multiple sequences), we analyzed a total of 4,168,042 non-redundant 16 amino acid long peptides.

**Figure 1.**
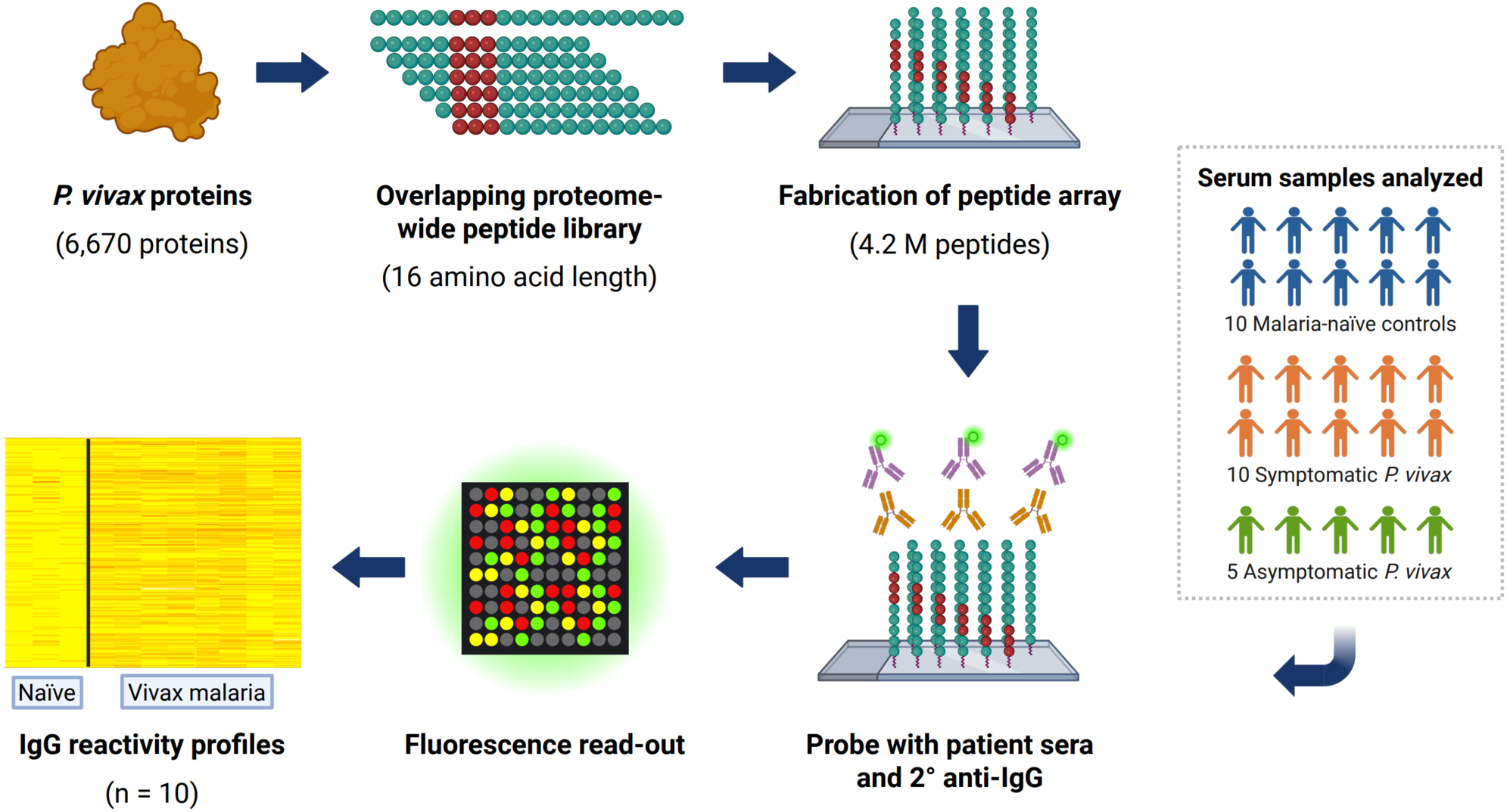
Summary of the peptide array design and data generation. Each *P. vivax* protein was bioinformatically split into every possible 16 amino acid peptide, and all non-redundant peptides were printed on a high-density array. Serum from one patient is then added to the array and IgG bound to a specific peptide detected using a fluorescently-labeled secondary antibody. Figure created with BioRender.

Microarrays were synthesized with a Nimble Therapeutics Maskless Array Synthesizer (MAS) by light-directed solid-phase peptide synthesis using an amino-functionalized support (Greiner Bio-One) coupled with a 6-aminohexanoic acid linker and amino acid derivatives carrying a photosensitive 2- (2-nitrophenyl) propyloxycarbonyl (NPPOC) protection group (Orgentis Chemicals). Amino acids (final concentration 20 mM) were pre-mixed for 10 min in N,N-Dimethylformamide (DMF, Sigma Aldrich) with N,N,N’,N’-Tetramethyl-O- (1H-benzotriazol-1-yl)uronium-hexafluorophosphate (HBTU, Protein Technologies, Inc.; final concentration 20 mM) as an activator, 6-Chloro-1-hydroxybenzotriazole (6-Cl-HOBt, Protein Technologies, Inc.; final concentration 20 mM) to suppress racemization, and N,N-Diisopropylethylamine (DIPEA, Sigma Aldrich; final concentration 31mM) as base. Activated amino acids were then coupled to the array surface for 3 min. Following each coupling step, the microarray was washed with N-methyl-2-pyrrolidone (NMP, VWR International), and site-specific cleavage of the NPPOC protection group was accomplished by irradiation of an image created by a Digital Micro-Mirror Device (Texas Instruments), projecting 365 nm wavelength light. Coupling cycles were repeated to synthesize the full in silico-generated peptide library.

Prior to sample binding, final removal of side-chain protecting groups was performed in 95% trifluoroacetic acid (TFA, Sigma Aldrich), 0.5% Triisopropylsilane (TIPS, TCI Chemicals) for 30 min. Arrays were incubated twice in methanol for 30 s and rinsed four times with reagent-grade water (Ricca Chemical Co.). Arrays were washed for 1 min in TBST (1× TBS, 0.05% Tween-20), washed 2× for 1 min in TBS, and exposed to a final wash for 30 s in reagent-grade water.

For each sample, 40 µl of serum were diluted 1:100 in binding buffer (0.01M Tris-Cl, pH 7.4, 1% alkali-soluble casein, 0.05% Tween-20) and bound to one array overnight at 4°C. After sample binding, the arrays were washed three times in wash buffer (1× TBS, 0.05% Tween-20), 10 min per wash. Binding of human IgG was detected via Alexa Fluor® 647-conjugated goat anti-human IgG secondary antibody (Jackson ImmunoResearch). The secondary antibody was diluted 1:10,000 (final concentration 0.1 ng/µl) in secondary binding buffer (1x TBS, 1% alkali-soluble casein, 0.05% Tween-20). Arrays were incubated with secondary antibody for 3 h at room temperature, then washed three times in wash buffer (10 min per wash), washed for 30 sec in reagent-grade water, and then dried by spinning in a microcentrifuge equipped with an array holder. Fluorescent signal of the secondary antibody was detected by scanning at 635 nm at 2 µm resolution using an Innopsys 1100AL microarray scanner (Innopsys Inc.). Scanned array images were analyzed with proprietary Nimble Therapeutics software to extract fluorescence intensity values for each peptide after (**Figure 1**).

### Determination of the serological response to P. vivax peptides

To identify *P. vivax* antigens among *P. vivax*-infected Cambodian adults, we used the analytical procedure summarized on **Figure S2**. We first considered, separately for each individual serum sample, whether the fluorescence intensity (FI) determined for a given peptide was greater than the FIs in all North American controls. To avoid including very low signals, we arbitrarily only considered peptides with FI greater than 500. Since each 16 amino acid peptide shares 15 amino acids with the peptides immediately preceding and following it in a given protein-coding sequence, we considered, as potentially antigenic, regions of proteins with at least seven consecutive peptides with signals greater than those of the malaria-naïve controls. The choice of using seven consecutive peptides (resulting in a shared examined region of 10 amino acids, **Figure S1**) was driven by the average length of reported linear B-cell epitopes (37, 38)) and not statistical considerations (statistical significance is rigorously determined later in the analyses, see below).

Finally, to allow for interindividual variability in the position of an antigenic region, we calculated the number of patients that were seropositive (*i.e.*, >=7 consecutive positive peptides) per 100 amino acid non-overlapping window across all proteins.

To rigorously assess statistical significance and identify antigenic proteins, we used a Poisson distribution to calculate the probability of observing by chance X patients seropositive for the same segment of 100 amino acid given the total number of potentially antigenic segments observed in one patient (**Supplemental Table 2**). Using this approach, we determined that the number of 100 amino acid segments identified as antigenic in seven or more of the ten patients was >20 fold greater than the number expected by chance alone (*i.e.*, corresponding to an FDR < 0.05) and we considered these segments as antigenic for the rest of the analyses.

### Intrinsically disordered regions and antigenic segments

To determine whether intrinsically disordered regions (IDRs) were disproportionally identified as antigenic on the peptide array, we first used the Predictors Of Natural Disordered Regions (PONDR) (48) to identify IDRs in all annotated *P. vivax* proteins using custom scripts. We then split each protein into the same 100 amino acid segments as described above and considered a segment as disordered if >50% of its amino acids were predicted as disordered (*i.e.*, the VL-XT score was greater than 0.5). We then compared whether disordered segments were enriched within antigenic segments using a χ-square test.

### Repeated peptides and antigenic segments

16 amino acid peptides present in multiple proteins, or multiple times in the same protein, were only printed once on the array. To determine if repeated peptides were disproportionally identified as antigenic, we compared whether segments containing >25% or >50% repeated peptides were enriched within the antigenic segments using a χ-square test.

### Amino acid composition of antigenic segments

To test if specific peptide motifs were disproportionally represented among antigenic segments, we calculated the number of all possible 5 amino acid peptides occurring in antigenic segments vs non-antigenic segments and compared their frequencies using Fisher’s exact test.

### Association between antigenicity and parasite mRNA expression in Cambodian individuals

To evaluate the association between the level of expression of a given protein and its antibody reactivity, we determined the average level of mRNA expression of all proteins in blood-stage parasites using bulk RNA-seq data generated from the same ten symptomatic *P. vivax* patients analyzed on the peptide array (49). Briefly, we reanalyzed the RNA-seq reads previously generated (BioProject: PRJNA378759) by mapping them with Hisat2 (50), first onto the human genome (NCBI Hg38 assembly) and then, for all the reads that did not align to the human genome, to the *P. vivax* genome (PlasmoDB-v68). After removing PCR duplicates using Picard, we calculated, for each patient, the number of reads mapped to each *P. vivax* gene. The average expression across all patients of each gene was then calculated after normalizing the counts obtained per patient to counts per million (CPM).

In order to determine if mRNA expression was statistically associated with antibody reactivity, we ranked all *P. vivax* genes tested on the peptide array based on their average mRNA expression and calculated, for 100 successive bins of ∼60 genes, the number of proteins that were immunoreactive in seven or more samples. We also tested whether the expression of a gene was statistically correlated with its seroreactivity using logistic regression.

## Results and Discussion

### Characterization of the seroreactivity of P. vivax proteins across symptomatic Cambodian individuals

We designed an ultra-dense peptide array containing 4.2 million peptides of 16 amino acids that cover the entire protein sequences of all *P. vivax* protein-coding genes (**Figures 1** and **S1**, see Materials and Method for details). We first probed these arrays with i) sera from ten vivax malaria Cambodian patients and ii) sera from ten malaria-naïve individuals and used a fluorescent secondary antibody against IgG to measure the patients’ antibody binding to each peptide. Out of the 4.2 million *P. vivax* peptides, 96,140 – 405,964 peptides per patient displayed a fluorescence intensity greater than the fluorescence measured in any of the ten malaria-naïve individuals and were further considered. Each 16 amino acid peptide on the array overlaps the next peptide in the protein sequence by 15 amino acids (**Figure S1**). Given that a typical linear B-cell epitope is on average 15 amino acids in length (38), we postulated that antibody binding to a specific protein should be detectable across multiple consecutive peptides. We therefore restricted our analyses to sections of proteins with, in a given patient, at least seven consecutive peptides with a signal greater than the signals observed in all malaria-naïve individuals (see Materials and Methods). We thus identified 3,332 – 16,197 putative immunoreactive regions per vivax malaria patient for a total of 81,362 putative immunoreactive regions across all patients (**Figure S2**). The fluorescence intensities detected in these putative immunoreactive regions were, quantitatively, relatively similar (typically with a signal intensity of 5,000-20,000). This observation contrasts with the different affinities and/or concentrations expected for natural antibodies targeting different antigens and likely indicated that the signal measured with the peptide array was only semiquantitative, either because of saturation of the fluorescence signal or data normalization. For the remaining analyses, we therefore exclusively focused on qualitative analyses and comparisons of the signals measured in infected individuals and controls.

We observed large variations in the sizes of the putative immunogenic regions, with a mean length of 10 amino acids (IQR=8-11) (**Figure S3**) but a few regions contained more than 100 consecutive putative immunoreactive peptides. Interestingly, the largest putative immunogenic region (142-145 amino acids long) was detected in the circumsporozoite protein (CSP, PVP01_0835600), one of the most studied *Plasmodium* antigenic proteins (51), and corresponded to a highly repetitive domain (see also below). Other proteins with very large immunoreactive domains included the surface protein P113 (52) (PVP01_1328200, 71-91 AA) and the asparagine-rich merozoite protein ARMA (53, 54) (PVP01_0937100, 99 AA). For many proteins, we observed several discontinuous putative antigenic regions (with an average of 5.7 regions identified in at least one patient per protein). One possible explanation for these multiple putative antigenic regions observed within one protein is that distant peptides (in the protein sequence) are in close proximity in the folded protein and represent a single 3D structure recognized by the host immune system. Alternatively, this observation could reflect that different antibodies specifically bind to different domains of the same protein.

While putative immunoreactive regions often overlapped among patients, many regions were not identified as seroreactive in all patients, or their exact boundaries differ between patients (see *e.g.*, **Figure S4**), suggesting that slightly different regions of the same protein might be immunogenic in different patients. Therefore, to identify proteins that were immunogenic across multiple patients while accounting for interindividual variations, we considered signals by non-overlapping 100 amino acid segments (see Materials and Methods). Finally, we evaluated statistical significance by comparing the number of 100 amino acid segments that were deemed seroreactive in X patients with the number expected by chance under a Poisson distribution and determined that segments that were immunoreactive in seven or more individuals were unlikely to be observed by chance only (FDR<0.05, **Supplemental Table 2**) and were considered antigenic. Overall, we identified a total of 315 immunogenic segments from 283 protein-coding genes (**Table 1**, **Supplemental Table 3**).

**Table 1:**
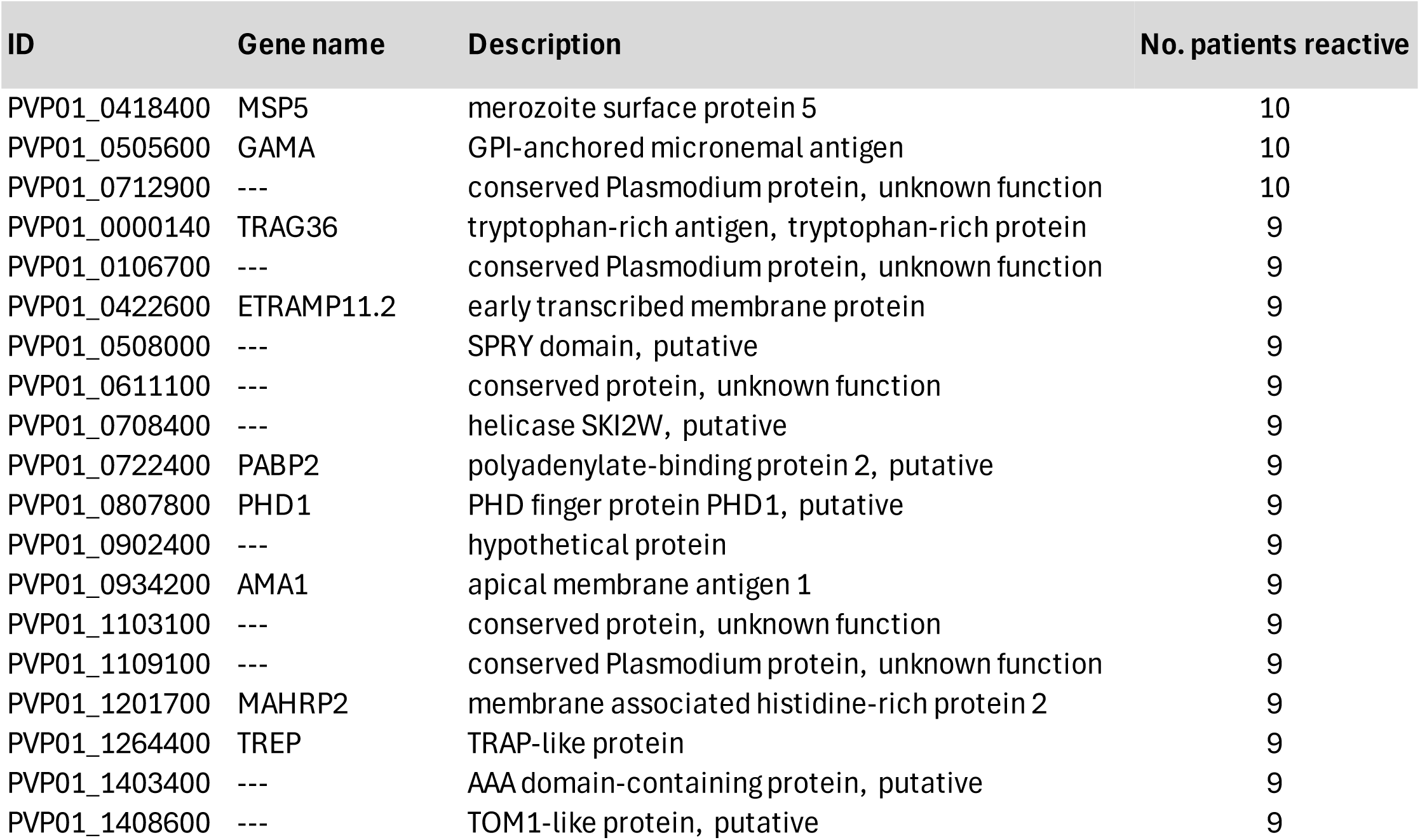
*P. vivax* proteins most commonly antigenic among symptomatic vivax malaria patients.

### General features of the immunogenic proteins detected

Both linear and conformational B-cell epitopes have been shown to play roles in protective immunity and immune evasion and are suitable for diagnostic purposes. For example, linear epitopes within the unstructured repeats of CSP are critical for protection (55–57), while some linear and/or repetitive asparagine-rich repeats may contribute to immune evasion (58–61). The high-density peptide array evaluates seroreactivity against linear epitopes and is limited in measuring the response to conformational epitopes (resulting from three-dimensional protein structures), unless a significant component of the conformational epitope is linear. To assess this limitation, we examined whether intrinsically disordered regions (IDRs) (62), which lack a predicted three-dimensional structure (but can sometimes change conformations based on interactions with other proteins or RNA molecules, were enriched among seroreactive segments. We observed that 37.3% of all immunoreactive segments) were disordered, a 1.4-fold enrichment compared with the proportion of disordered regions in non-reactive segments (p = 2.2x10^-16^). Note however that 62.7% of the detected immunoreactive segments were derived from regions of the proteins with predicted tertiary structure, indicating that the immunogenic segments identified were not solely restricted to disordered domains.

Next, we evaluated whether repeated peptides (*i.e.*, peptides printed once on the array but present multiple times in one protein or in multiple proteins) were more likely to be seroreactive than were unique peptides. Among the 315 immunoreactive segments, only 10 (3.2%) contained >25% repeated peptides, very similar to the proportion among the non-seroreactive peptides (4.4%, p = 0.3), indicating that repeated peptides were not significantly more seroreactive in our dataset (similar results were obtained when considering segments with >50% repeated peptides).

We then examined whether specific amino acid motifs were over-represented in the immunoreactive segments. Three 5-mers, present in 17 different immunogenic proteins, were significantly enriched in immunoreactive segments: ANAAN, NAANA and AANAA (p = 3.59 x 10^-40^, p = 1.48 x 10^-26^, p = 3.39 x 10^-22^ respectively, Fisher’s exact test, **Figure S5**). Interestingly, asparagine- and alanine-rich repeats have been reported in several *P. falciparum* antigens, possibly as “decoy epitopes” triggering a highly antigenic but non-protective immune response (61, 63, 64).

Finally, we examined whether the immunogenicity of a protein was associated with its level of expression. Since we lack comprehensive quantitative data on protein abundance, we used the mRNA expression levels determined by whole-blood RNA-seq generated from the same symptomatic *P. vivax* infections as a proxy (49). While the level of mRNA expression determined from patient blood is influenced by the stage composition of each infection, we noted that, overall, peptides derived from proteins that were highly expressed during a blood infection were more likely to be seroreactive (**Figure 2**, p = 1.28 x 10^-15^), suggesting that the amount of a protein present during a blood stage infection may be an important factor influencing the ability of the immune system to mount a humoral immune response.

**Figure 2.**
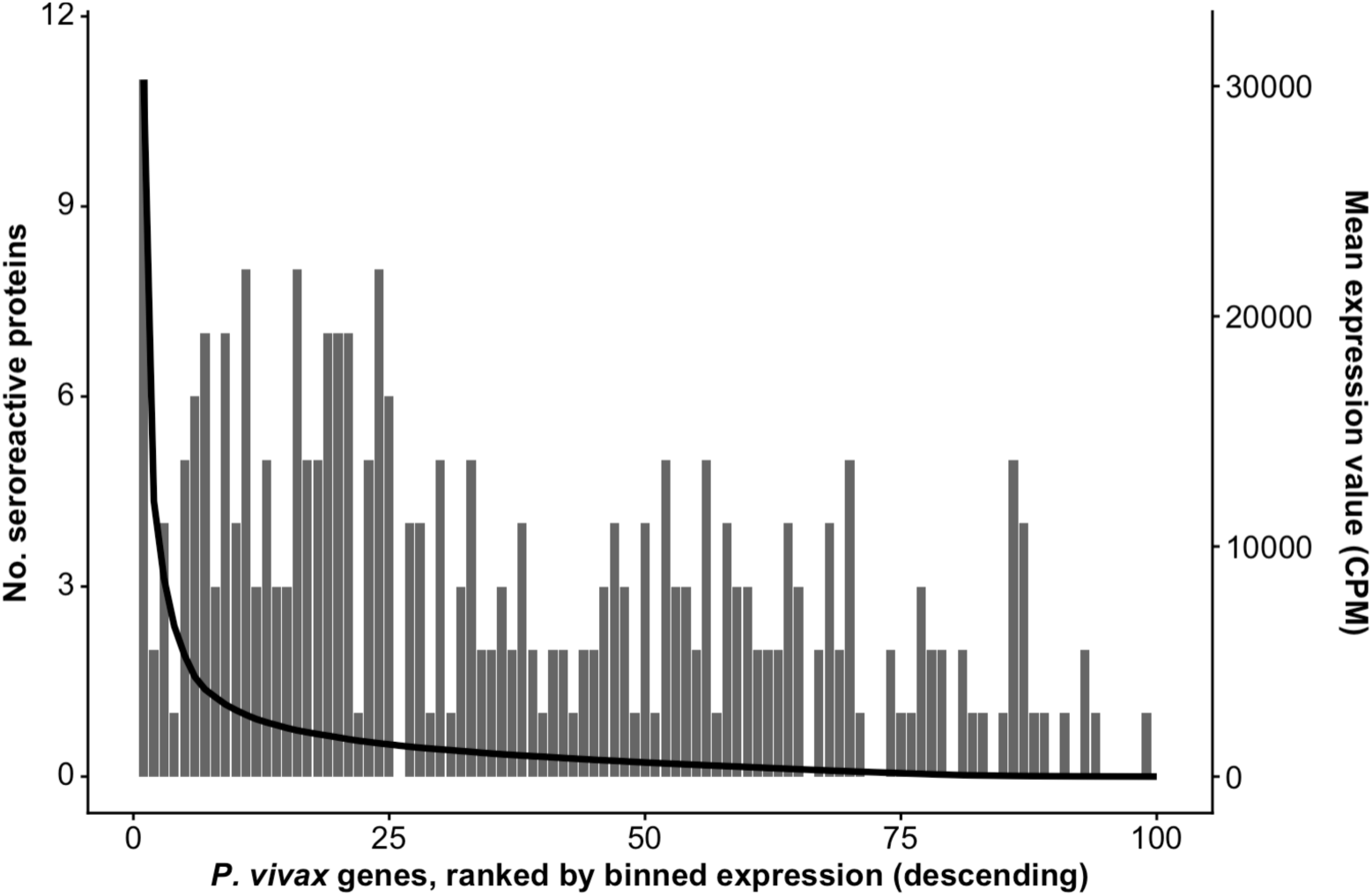
Association between the level of mRNA expression of a gene in blood stage parasites and its protein antigenicity. The x-axis shows 100 bins of 66 *P. vivax* genes grouped according to their average mRNA expression in blood stage parasites (black line, right y-axis) and ranked from left (most expressed) to right (least expressed). The vertical bars indicate the number of antigenic proteins in each bin (right y-axis).

### Many erythrocyte invasion proteins are highly immunogenic

Among the 283 immunogenic *P. vivax* proteins, many proteins known to be involved in erythrocyte invasion were detected (**Figure 3**). For example, Merozoite Surface Protein 5 (MSP5), Tryptophane-Rich Antigen 36 (TRAg36) and Apical Membrane Antigen 1 (AMA1) are among the most common antigenic proteins (**Table 1**).

**Figure 3.**
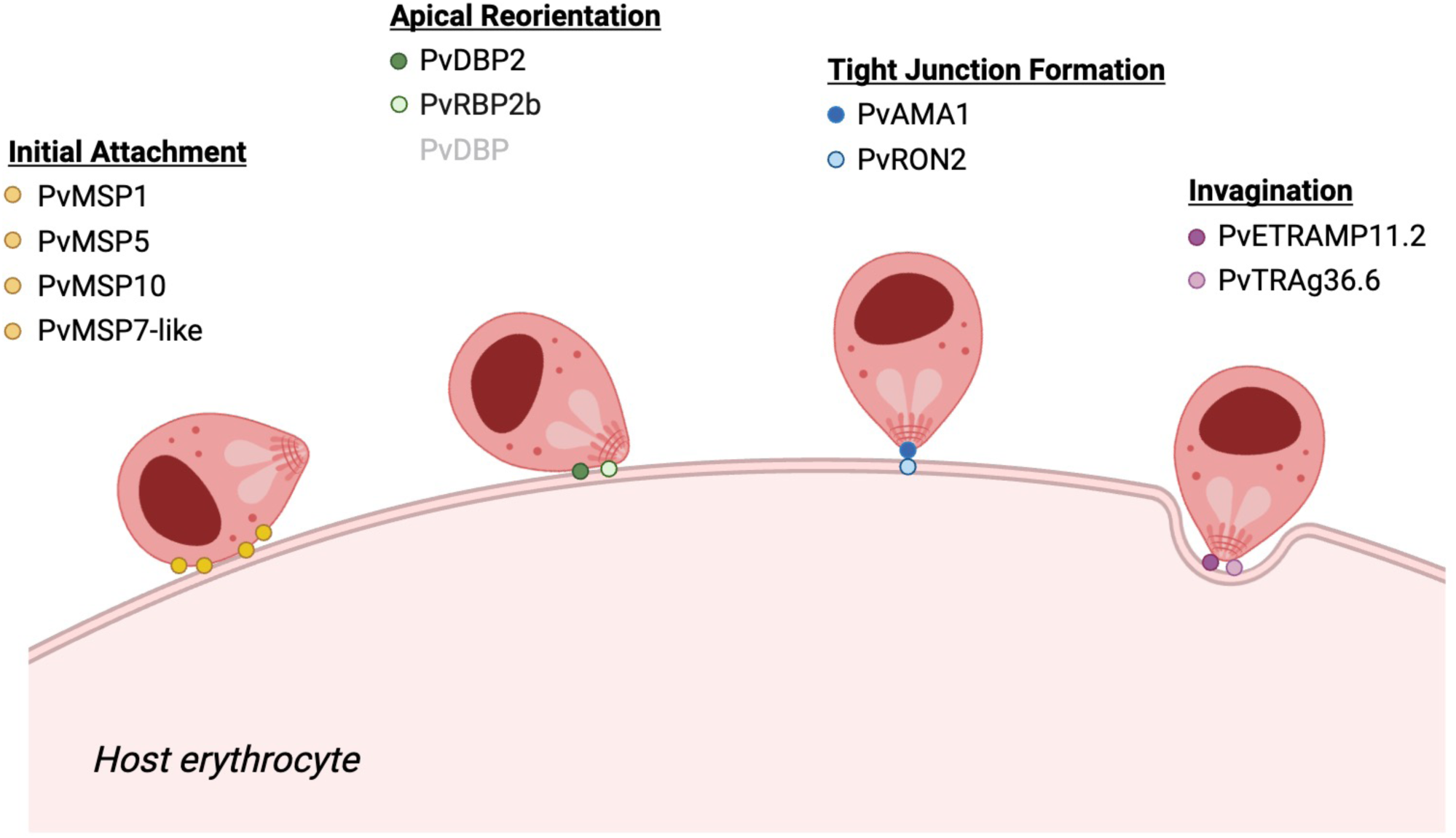
*P. vivax* proteins involved in erythrocyte invasion are highly immunogenic. The figure shows the proteins known to be involved at different stages of red blood cell invasion. Proteins deemed immunogenic on our high-density peptide array are indicated in black while proteins under the significance threshold are in grey. Figure was created with BioRender.

Antibodies against MSP5 (PVP01_0418400) were detected in all ten patients, whereas antibodies against MSP10 (PVP01_1129100), MSP1 (PVP01_0728900), and MSP7-like protein (PVP01_1220300) were detected in more than seven patients (**Supplemental Table 3, Figure S6**). Merozoite surface proteins are “adhesins” involved in the initial attachment of the parasite to erythrocytes prior to invasion (65, 66). Most of these proteins remain incompletely characterized, especially in *P. vivax,* but might act as part of complexes with other MSPs and additional proteins and are already present on the surface of the merozoites upon egress from the schizonts. Only MSP1 has been the focus of detailed serological studies in *P. vivax,* and the C-terminal domain (MSP1-42 kDa) has notably been shown to be highly antigenic (67–69), with antibodies against this region being detectable several months after one infection (70). The signal detected on the peptide array overlapped with the 42 kDa region but appeared to be restricted to the highly polymorphic MSP1-33 kDa fragment rather than located in the MSP1-19 kDa region, the putative RBC binding domain and a major target of protective immunity (71) (**Figure 4A**). Peptides from the apical membrane antigen 1 (AMA1, PVP01_0934200) were recognized in nine of the ten patients and displayed antibody binding signal in the C-terminal region that overlapped with the cytoplasmic tail (CT) domain (72, 73) (**Figure 4B**). AMA1 is a micronemal protein found on the apical end of merozoites that mediates the initiation of tight junction formation (74, 75) (**Figure 3**). While it is only secreted from the microneme at the time of invasion (76), AMA1 is one of the best characterized *Plasmodium* antigens and is a leading candidate for monoclonal antibody therapy (77–79) and vaccine design (80). Similar to MSP1, the location of antigenic signal detected by the peptide array was located in the C-terminal domain. In both *P. falciparum* and *P. vivax*, the 22 kDa cytoplasmic tail is carried into the host cell after cleavage of the AMA1 ectodomain, which is thought to facilitate evasion of protective anti-AMA1 immune responses, primarily targeting domain II (81) (**Figure 4B**). The binding partner of AMA1, the rhoptry neck protein 2 (RON2, PVP01_1255000) was also identified as immunogenic, with an antibody binding signal located at the N-terminus of the protein (**Figure S7**). RON2 is a parasite protein that is stored in the rhoptry neck and injected into erythrocytes, before becoming embedded in the host membrane, where it serves as a ligand for AMA1 to initiate the tight junction formation and trigger erythrocyte invasion (76, 82).

**Figure 4.**
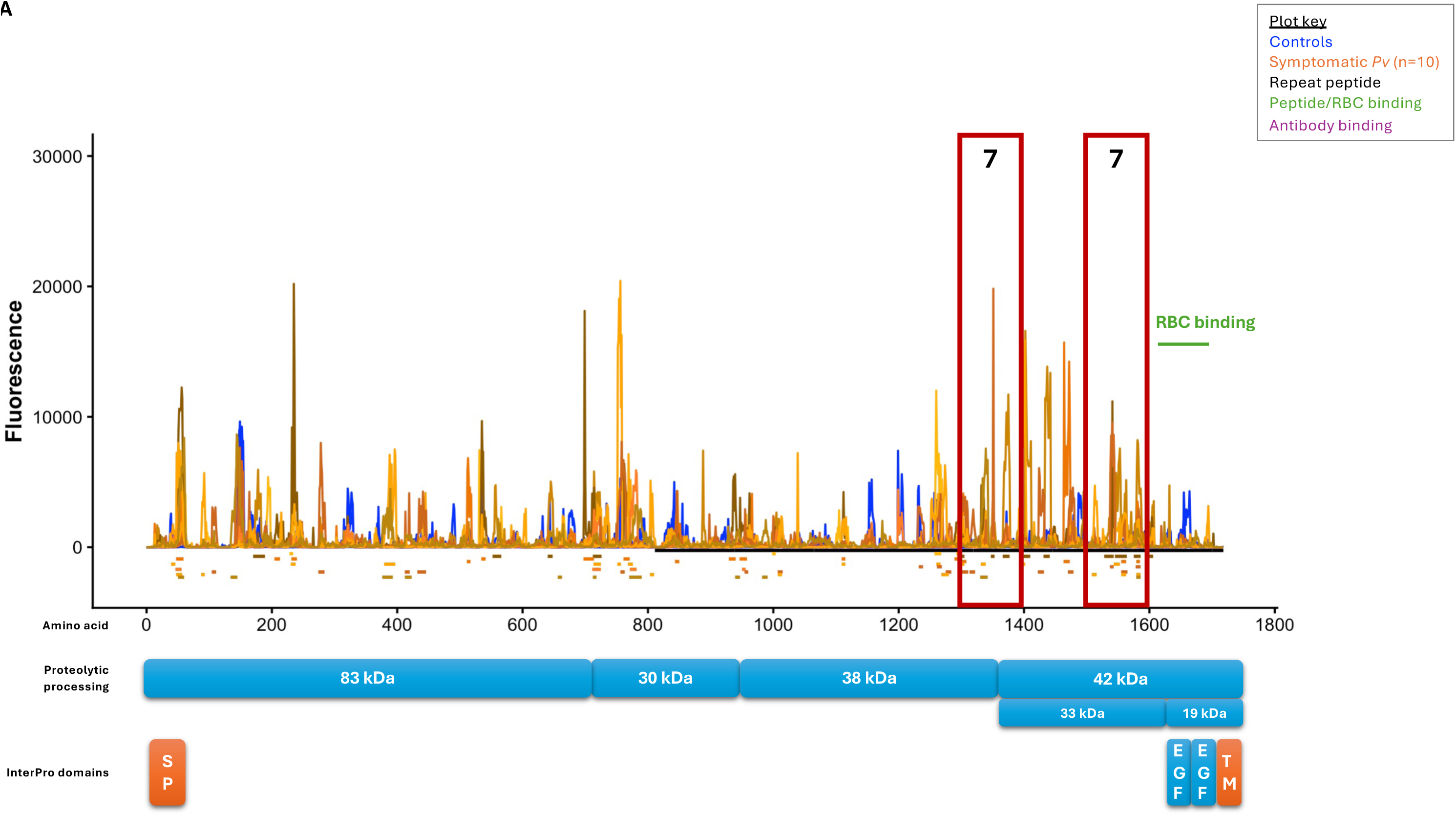
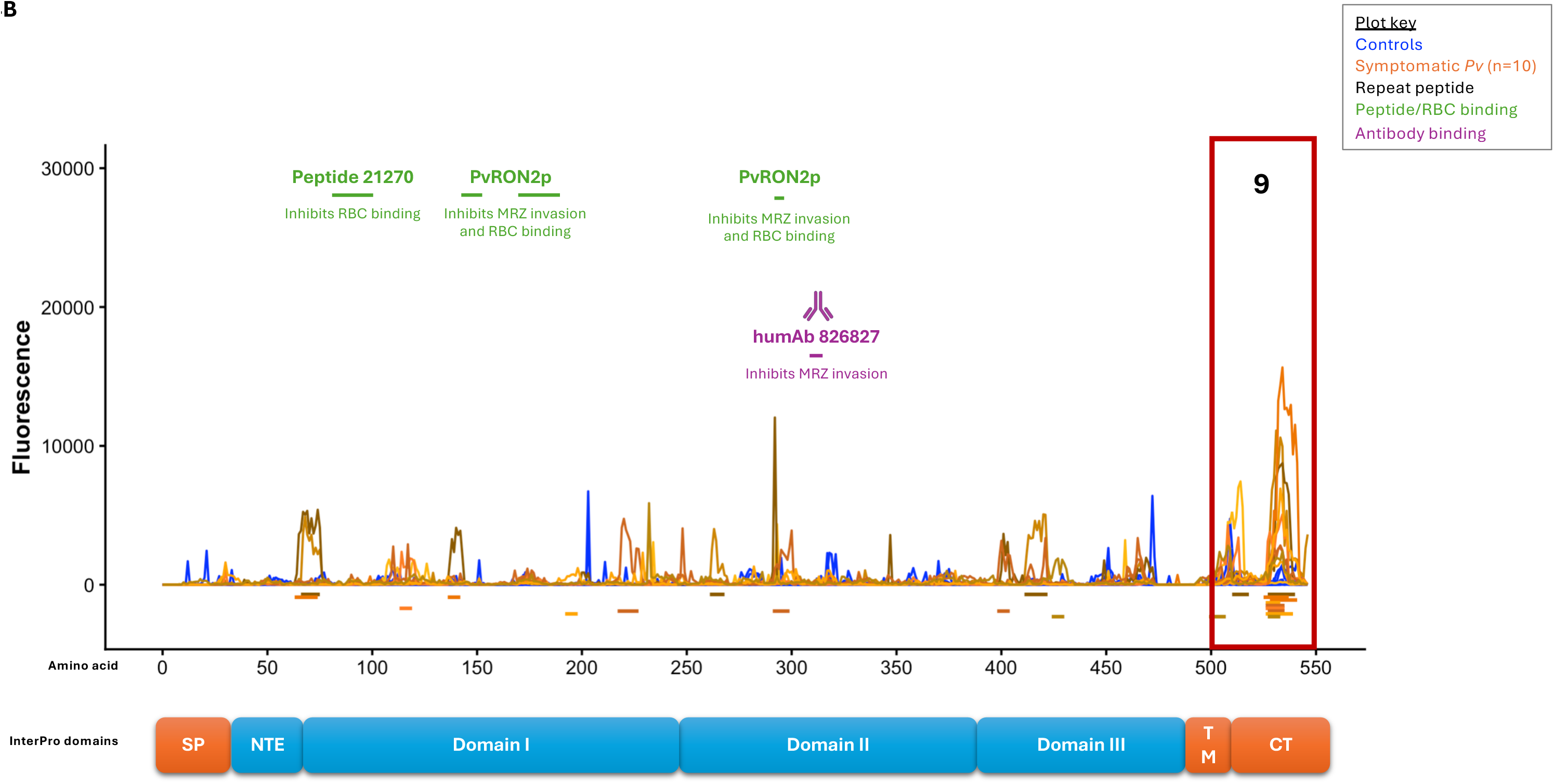
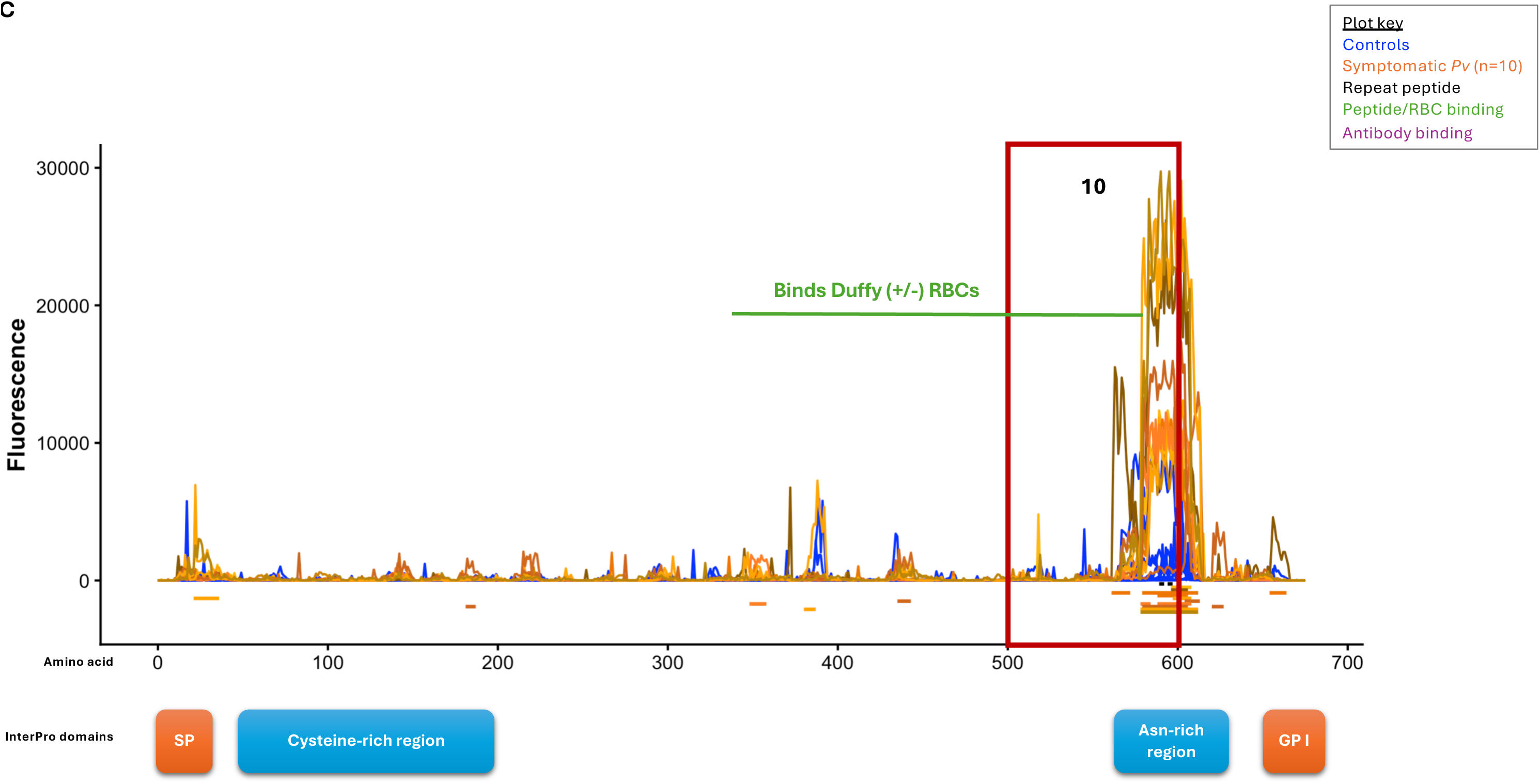
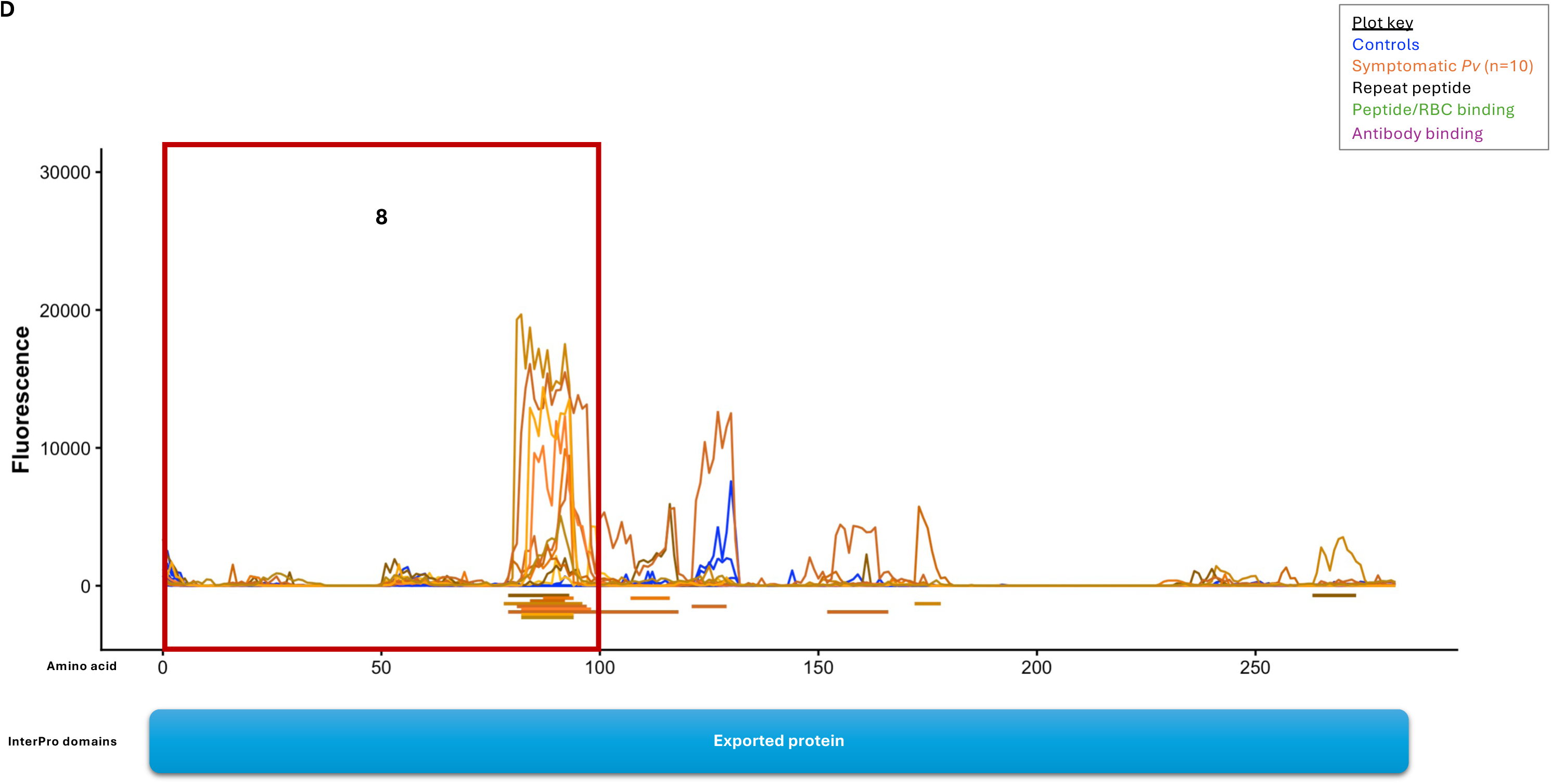
IgG antibody binding across four *P. vivax* proteins: **(A)** Merozoite surface protein 1 (MSP1, PVP01_0728900), **(B)** Apical membrane antigen 1 (AMA1, PVP01_0934200), **(C)** GPI-anchored micronemal protein (GAMA, PVP01_0505600), **(D)** *Plasmodium* exported protein (PVP01_1201600). Each panel shows the fluorescence detected (y-axis) at peptides organized according to the protein amino acid sequence (x-axis). The blue lines show the data for each malaria-naïve control and the yellow to red lines represent the symptomatic individuals (n=10). Each red box indicates an antigenic 100 amino acid segment. The blue and orange boxes under each plot indicate annotated functional regions and/or proteolytic domains. The green bars indicate regions of the protein known to be associated with RBC bindings, while the purple antibody indicates a human monoclonal antibody inhibiting RBC invasion.

Several tryptophan-rich antigen (TRAg) genes are also among the most common seroreactive proteins, including TRAg36 (PVP01_0000140) and TRAg38 (PVP01_0503600), which may bind to Band 3 on the host erythrocyte (83, 84), as well as TRAg32/36.6 (PVP01_0000170) and its binding partner ETRAMP11.2 (PVP01_0422600) (85) (**Figure S8).** TRAgs are members of a multigene family present in all *Plasmodium* species but are significantly expanded in *Plasmodium vivax* (*86*). While the exact role of these proteins remains unclear, several of these proteins are able to bind to RBCs, possibly via lipid binding (87) or directly to host proteins (84, 88), and have been shown to be antigenic (86).

Finally, we identified the Duffy binding protein 2 (DBP2, PVP01_0102300) (89), also referred to as erythrocyte binding protein (EBP or EBP2), among the most immunogenic proteins (**Supplemental Table 3, Figure S9A**). DBP2 may mediate a Duffy-independent invasion pathway (90), possibly through binding to complement receptor 1 (91). The Duffy-binding protein (DBP, PVP01_0623800), one of the most studied *P. vivax* proteins and a critical reticulocyte invasion protein, did not reach significance in our analyses (**Figure S9B**). This lack of signal contrasts with previous studies that have clearly demonstrated humoral immunity against DBP (92–94) but may be explained by the main limitation of peptide arrays, which primarily identify linear epitopes: several studies have shown that antibodies to DBP bind mainly conformational epitopes (95–97) and are therefore unlikely to be captured by short amino acid peptides. Similarly, we failed to detect antibody binding signals for most Reticulocyte Binding Proteins (RBPs (98)), with the exception of RBP2b (PVP01_0800700, **Figure S10**), which contrasts with the results of previous serological studies but may be explained by the conformational epitopes of these proteins.

Overall, the identification of most key invasion genes among seroreactive *P. vivax* proteins is consistent with the localization of these proteins on the surface of the merozoites and, therefore, their exposure to the host immune system (while most *P. vivax* proteins are hidden inside the parasites and/or the RBCs and are less likely to be recognized). While protective antibodies have been reported for several of these invasion proteins, it is noteworthy that the signal of antibody binding detected among the Cambodian patients typically did not overlap with these protective antibody binding regions. One possible explanation for this observation is that the individuals analyzed here were experiencing a symptomatic episode of vivax malaria and, therefore, had not developed protective immunity against the parasites (but also see below) and that the signal identified reflected the humoral detection of these immunogenic proteins but, possibly, from domains implicated in immune evasion.

We also detected antibody-binding signal for one putative invasion protein that has received little attention in *P. vivax*: the GPI-anchored micronemal protein (GAMA, PVP01_0505600). Peptides of the protein were recognized by antibodies from all ten patients, with a clearly defined peak of reactivity centered on the asparagine-rich region at the C-terminus of the protein (**Table 1**, **Figure 4C**). While this protein has not been extensively studied in *P. vivax,* it is believed to play a role in reticulocyte invasion, possibly via interactions with MSP10 (99). The observation of shared seroreactivity for GAMA, suggests that it is accessible by the host immune system, similar to many known erythrocyte invasion proteins, and may indicate that it plays a key role in this process in *P. vivax*, warranting further investigations.

### SEC13 and other nucleoporins are enriched among immunogenic proteins

A second group of proteins, nucleoporins, which form pore complexes on the nuclear envelope of *Plasmodium* parasites (100), were significantly enriched among the proteins eliciting a humoral immune response (p = 1.37 x 10^-6^). We identified six nucleoporins that were immunogenic among Cambodian patients: NUP313, NUP205, NUP138, NUP434, SEC13, and SEC31 (**Supplemental Table 3, Figure S11**). This finding was somewhat surprising, as in contrast to invasion proteins that are displayed at the surface of the merozoites and therefore exposed to the host immune system, nucleoporins are believed to be hidden within the parasite, behind the RBC- and the parasitophorous vacuolar membranes. However, these proteins may also participate in other functions, with different cellular localizations that may expose them to the host immune system. Notably, SEC13 seems to be localized, at least in *P. falciparum* gametocytes and sporozoites, outside the nucleus and may be associated with proteins involved in secretion or invasion mechanisms (101). Our data would be consistent with such a dual role and support further investigations of the function of these proteins.

### Some P. vivax hypothetical proteins are highly immunogenic

Half of the *P. vivax* genes encode proteins without functional annotation. Among the 283 highly immunogenic proteins, 104 fell within this category and were labeled solely as “conserved”, “hypothetical proteins” or “exported proteins” without further annotations (**Supplemental Table 3**). The identification of antibody-binding signals for these 104 proteins could indicate that they are located on the surface of merozoites, exported to the surface of the RBCs, or possibly secreted. One interesting example is PVP01_1201600, which is only annotated as an exported protein. This protein is only 298 amino acids long but displays an antibody binding signal localized between the 75^th^ and 100^th^ amino acids and identified in eight out of the ten patients (**Table 1**, **Figure 4D**). This gene has no ortholog in *P. falciparum* but is located, in *P. vivax*, immediately upstream of HRP2 (PVP01_1201700, a highly abundant secreted protein used in *Plasmodium* RDTs) and is the 28^th^ most highly expressed gene by blood-stage parasites. This immunogenicity, combined with this high expression pattern, is fascinating and points to a unique role for this short protein during blood-stage infections and warrants further investigations. Similarly, we observed similar antibody-binding signals in PVP01_1103100, PVP01_1426900, PVP01_1271400, and PVP01_1030800 (**Figure S12**), suggesting that these proteins may be particularly exposed to the host immune system and that their examination should be prioritized in functional studies.

### Low frequency of antibody binding to sporozoite peptides

The circumsporozoite protein (CSP, PVP01_0835600), which covers the entire surface of sporozoites (51) and is the target of the two current malaria vaccines in *P. falciparum* (RTS,S and R21 (102)), exhibited the longest stretch of seroreactive amino acids. However, CSP was recognized by antibodies in only five out of ten patients and therefore did not reach our significance threshold (**Table S3, Figure S13**). This lack of antibody binding in some patients, especially compared with the signal identified for many erythrocyte invasion proteins, could be due to the diversity of CSP alleles present in the *P. vivax* population: at least three different alleles, VK210, VK247, and *P. vivax*-like, are common among *P. vivax* parasites, with extensive amino acid differences among them, especially in the central repeat region (amino acid range 97 – 275) (51, 103). Unfortunately, only peptides corresponding to CSP-VK210 were included in the peptide array. The antibody binding signal detected at CSP is located between amino acid positions 240 and 275, in the repetitive and highly divergent region (**Figure S13**); therefore, the lack of signal in five patients could be due to exposure to one of the other alleles that was not captured in our data but circulates in the Cambodian *P. vivax* parasite population (104). Alternatively, this low frequency of anti-CSP antibodies could reflect the fact that many infections caused by *P. vivax* derived from the reactivation of hypnozoites and not directly from an infectious bite (105) and that some of the patients studied might not have been recently exposed to sporozoites.

To evaluate this latter possibility, we examined antibody binding to other key sporozoite proteins. TREP (PVP01_1264400) was the only sporozoite protein that reached the significance threshold, and for which antibodies were detected in nine of the ten patients (**Figure S13**). In rodent parasites, the TREP protein is involved in the gliding motility of sporozoites and might therefore be exposed to, and recognized by, the host immune system (106). In contrast, antibodies against TRAP (PVP01_1218700), another pre-erythrocytic vaccine candidate evaluated alongside CSP (107, 108), and MAEBL (PVP01_0948400), a sporozoite adhesin essential notably for attachment to the mosquito salivary glands (109, 110), were detected in only six and five patients, respectively (**Figure S13**). Overall, antibody binding to sporozoite proteins reaches significance less frequently than binding to peptides derived from proteins involved in erythrocyte invasion, consistent with the low immunogenicity of pre-erythrocytic stages and possibly reflecting the major differences in the number of parasites from these stages. Antibodies from two of the ten patients did not recognize either CSP, TRAP or MAEBL (but were positive for TREP). One could speculate that these patients’ infections were relapses caused by the reactivation of dormant liver-stage parasites and that these individuals may not have been exposed to an infectious bite for several months, explaining their absence of detectable antibody reactivity to sporozoite proteins. Additional analyses, including a larger cohort of patients and longitudinal data to evaluate antibody decay over time, will be necessary to determine whether antibody reactivity to these proteins could be informative for discriminating relapses from new infections. It would also be interesting to robustly characterize the expression (or lack thereof) of TREP in sporozoites and blood-stage *P. vivax* parasites, as, if TREP is indeed exclusively expressed in sporozoites, measurement of seroreactivity against TREP may allow for more robust detection of infective bites and assessment of ongoing transmission.

### The antibody binding profiles of asymptomatic vivax malaria individuals differ from those of patients

In addition to symptomatic patients, we also probed the peptide array with sera from five asymptomatic Cambodian individuals infected with *P. vivax.* The smaller sample size prevented us from rigorously assessing the statistical significance and determining with confidence which proteins elicited a humoral immune response in these individuals. Nonetheless, we observed 483 *P. vivax* regions (from 414 proteins) whose fluorescence signal was greater than that of all malaria-naïve individuals in all five asymptomatic individuals tested (**Supplementary Table 4**). Interestingly, this number is 5-fold greater than the average number of regions significantly immunoreactive in five random symptomatic individuals: when we randomly sampled five patients out of the ten symptomatic individuals screened on the peptide array, we identified only 29 - 146 regions (with a mean of 90) that were significant in all of them (compared with the 483 regions observed in five asymptomatic individuals). While this finding needs to be validated in a much larger cohort, it could indicate that asymptomatic individuals have antibodies against a broader repertoire of *P. vivax* proteins than symptomatic individuals. One possible explanation for this observation is that this broader repertoire of antibodies against *P. vivax* proteins may help individuals control the infection and prevent the development of malaria symptoms, although it is important to reiterate here that the peptide array does not provide any information on the protective nature of the antibodies detected. Indeed, examination of the profile of antibody binding along the MSP1, AMA1 or DBP protein sequences failed to reveal a significant signal overlapping the binding location of known protective antibodies (**Figure S14**). An alternative, but not exclusive, explanation for the larger number of recognized proteins in asymptomatic individuals is that it may result from longer exposure to malaria parasites and the development of a broader, but not necessarily protective, antibody response: while we do not have any information regarding the time of infection of any of the individuals, it is likely that asymptomatic individuals have chronic and long-sustained infections, whereas the symptomatic individuals may result from a recent, uncontrolled infection. Further studies, including longitudinal monitoring of the antibody responses of both asymptomatic and symptomatic individuals, will be necessary to differentiate between these hypotheses.

The small sample size of the asymptomatic cohort analyzed here prevented us from rigorously determining which specific proteins were statistically seroreactive in asymptomatic individuals. However, among the 414 proteins that elicited antibody responses in all five asymptomatic individuals, we observed 69 PIR genes. PIR genes are members of a highly variable multigene family that are possibly responsible for antigenic variation in *P. vivax* (111). Out of the 69 PIR segments that were immunogenic in all asymptomatic individuals, only seven were deemed antigenic in symptomatic patients (*i.e.*, detected in seven or more patients). In fact, we detected antibodies against twice as many PIR proteins, proportionally, in asymptomatic individuals than in symptomatic patients (15.7% vs 7.1%, p=0.0006). While this result needs to be confirmed, the increased frequency of antibodies elicited against PIR proteins observed in asymptomatic patients could suggest that these antibodies play a protective role against clinical malaria.

## Conclusion

We identified 283 proteins that were immunogenic among symptomatic vivax patients and that included most of the proteins known to be involved in RBC invasion. While the domains identified as immunogenic in our study differed from domains previously reported to be associated with protective immunity in DBP, AMA1 or MSP1, immunogenic proteins identified here could provide new targets for vaccine development. In particular, we observed immunoreactivity of a putative invasion protein, several nucleoporins and uncharacterized proteins that should be further investigated for their roles during blood-stage infections. Our analyses also revealed a unique binding pattern of antibodies targeting PIR proteins in asymptomatic individuals, which may be associated with protection against clinical vivax malaria. Overall, the analyses presented here provide an agnostic and comprehensive resource for the malaria community to develop new tools for the detection and elimination of this important human pathogen.

## Acknowledgements

We thank all patients and health care workers involved in this study and the staff of the Malaria Research Unit at the Institut Pasteur of Cambodia and the staff of the National Center for Parasitology, Entomology and Malaria Control in Cambodia for their collaboration and sample collection. This study was supported by awards from the NIH to DS (R21AI159307 and R01AI153083). JP is supported by the NIH/NIAID (R01AI173171, R01AI175134 and R61AI187100) and by the Pasteur International Unit PvESMEE. The funders had no role in study design, data collection and analysis, decision to publish, or preparation of the manuscript. This research was supported by the Intramural Research Program of the National Institutes of Health (NIH). The contributions of the NIH author(s) are considered Works of the United States Government. The findings and conclusions presented in this paper are those of the author(s) and do not necessarily reflect the views of the NIH or the U.S. Department of Health and Human Services.

## Data and code availability

All data generated in this study is available at https://doi.org/10.5281/zenodo.18853563. Custom scripts are available at https://github.com/rosita-asawa.

## Figure legends

**Figure S1.** Schematic showing consecutive peptides of 16 amino acid (green horizontal bars) overlapping by 15 amino acids. Analysis of seven consecutive peptides leads to a shared region of 10 amino acids (purple box).

**Figure S2**. Overview of the analytical pipeline used to determine IgG antibody binding within the peptide array.

**Figure S3**. Distribution of the number of consecutive seroreactive peptides detected on the peptide array. Note that the x axis is shortened for display purposes.

**Figure S4**. Example of antibody binding against one protein (MSP5, PVP01_0418400) illustrating extensive inter-individual variations in antibody binding among patients. The x-axis represents the position of consecutive peptides assessed by the high-density array and the y-axis the fluorescence measured at each peptide. The yellow, orange and brown lines show the IgG antibody binding for three Cambodian patients, while the blue lines represent data from all malaria-naïve individuals (n=10). The horizontal lines under the plot show locations where seven or more consecutive peptides have a signal greater than all malaria-naïve controls (colored by patient). The blue and orange boxes under each plot indicate annotated functional regions.

**Figure S5**. Distribution of all 5 amino acid motifs across antigenic segments (y-axis) and non-antigenic segments (x-axis).

**Figure S6**. Antigenic profiles of merozoite surface proteins (MSPs). See legend of Figure 4 for details.

**Figure S7**. Antigenic profile across the rhoptry neck protein 2 (RON2, PVP01_1255000). See legend of Figure 4 for details.

**Figure S8**. Antigenic profiles of tryptophan-rich antigens (TRAgs). See legend of Figure 4 for details.

**Figure S9. (A)** Antigenic profile across the Duffy binding protein 2 (DBP2, PVP01_0102300). **(B)** Antigenic profile across the Duffy binding protein (DBP, PVP01_0623800). See legend of Figure 4 for details.

**Figure S10**. Antigenic profiles of reticulocyte binding proteins. See legend of Figure 4 for details.

**Figure S11**. Antigenic profiles of nucleoporins. See legend of Figure 4 for details.

**Figure S12**. Antigenic profiles across selected hypothetical proteins. See legend of Figure 4 for details.

**Figure S13**. Antigenic profiles across selected sporozoite-related proteins. See legend of Figure 4 for details.

**Figure S14**. Antigenic profiles of the asymptomatic individuals across selected invasion proteins with known protective antibodies. See legend of Figure 4 for details.

## References

1. World Health Organization. 2025. World Malaria Report 2025. World Health Organization, Geneva.

2. Anstey NM, Tham WH, Shanks GD, Poespoprodjo JR, Russell BM, Kho S. 2024. The biology and pathogenesis of vivax malaria. Trends Parasitol 40:573–590.

3. Adams JH, Mueller I. 2017. The Biology of Plasmodium vivax. Cold Spring Harb Perspect Med 7.

4. Malleret B, Li A, Zhang R, Tan KSW, Suwanarusk R, Claser C, Cho JS, Koh EGL, Chu CS, Pukrittayakamee S, Ng ML, Ginhoux F, Ng LG, Lim CT, Nosten F, Snounou G, Rénia L, Russell B. 2015. Plasmodium vivax: restricted tropism and rapid remodeling of CD71-positive reticulocytes. Blood 125:1314–1324.

5. Kanjee U, Rangel GW, Clark MA, Duraisingh MT. 2018. Molecular and cellular interactions defining the tropism of Plasmodium vivax for reticulocytes. Curr Opin Microbiol 46:109–115.

6. Groves ES, Simpson JA, Edler P, Daher A, Pasaribu AP, Pereira DB, Saravu K, von Seidlein L, Rajasekhar M, Price RN, Commons RJ, WorldWide Antimalarial Resistance Network PvFSG. 2025. Parasitaemia and fever in uncomplicated Plasmodium vivax malaria: A systematic review and individual patient data meta-analysis. PLoS Negl Trop Dis 19:e0012951.

7. White NJ. 2011. Determinants of relapse periodicity in Plasmodium vivax malaria. Malar J 10:297.

8. Battle KE, Karhunen MS, Bhatt S, Gething PW, Howes RE, Golding N, Van Boeckel TP, Messina JP, Shanks GD, Smith DL, Baird JK, Hay SI. 2014. Geographical variation in Plasmodium vivax relapse. Malaria Journal 13:144.

9. Chu CS, White NJ. 2021. The prevention and treatment of Plasmodium vivax malaria. PLOS Medicine 18:e1003561.

10. Mueller I, Galinski MR, Baird JK, Carlton JM, Kochar DK, Alonso PL, del Portillo HA. 2009. Key gaps in the knowledge of Plasmodium vivax, a neglected human malaria parasite. Lancet Infect Dis 9:555–66.

11. Eng V, Lek D, Sin S, Feufack-Donfack LB, Orban A, Salvador J, Seng D, Heng S, Khim N, Tebben K, Flamand C, Sommen C, van der Pluijm RW, White M, Witkowski B, Serre D, Popovici J. 2025. 14 days of high-dose versus low-dose primaquine treatment in patients with Plasmodium vivax infection in Cambodia: a randomised, single-centre, open-label efficacy study. Lancet Infect Dis 25:884–895.

12. Nkhoma ET, Poole C, Vannappagari V, Hall SA, Beutler E. 2009. The global prevalence of glucose-6-phosphate dehydrogenase deficiency: a systematic review and meta-analysis. Blood Cells Mol Dis 42:267–278.

13. Duffy PE, Gorres JP, Healy SA, Fried M. 2024. Malaria vaccines: a new era of prevention and control. Nature Reviews Microbiology 22:756–772.

14. Dickey TH, Tolia NH. 2023. Designing an effective malaria vaccine targeting Plasmodium vivax Duffy-binding protein. Trends Parasitol 39:850–858.

15. Draper SJ, Sack BK, King CR, Nielsen CM, Rayner JC, Higgins MK, Long CA, Seder RA. 2018. Malaria Vaccines: Recent Advances and New Horizons. Cell Host Microbe 24:43–56.

16. Arévalo-Herrera M, Gaitán X, Larmat-Delgado M, Caicedo MA, Herrera SM, Henao-Giraldo J, Castellanos A, Devaud J-C, Pannatier A, Oñate J, Corradin G, Herrera S. 2022. Randomized clinical trial to assess the protective efficacy of a Plasmodium vivax CS synthetic vaccine. Nature Communications 13:1603.

17. Bennett JW, Yadava A, Tosh D, Sattabongkot J, Komisar J, Ware LA, McCarthy WF, Cowden JJ, Regules J, Spring MD, Paolino K, Hartzell JD, Cummings JF, Richie TL, Lumsden J, Kamau E, Murphy J, Lee C, Parekh F, Birkett A, Cohen J, Ballou WR, Polhemus ME, Vanloubbeeck YF, Vekemans J, Ockenhouse CF. 2016. Phase 1/2a Trial of Plasmodium vivax Malaria Vaccine Candidate VMP001/AS01B in Malaria-Naive Adults: Safety, Immunogenicity, and Efficacy. PLOS Neglected Tropical Diseases 10:e0004423.

18. Herrera S, Fernández OL, Vera O, Cárdenas W, Ramírez O, Palacios R, Chen-Mok M, Corradin G, Arévalo-Herrera M. 2011. Phase I Safety and Immunogenicity Trial of Plasmodium vivax CS Derived Long Synthetic Peptides Adjuvanted with Montanide ISA 720 or Montanide ISA 51. The American Society of Tropical Medicine and Hygiene 84:12–20.

19. Hou MM, Barrett JR, Themistocleous Y, Rawlinson TA, Diouf A, Martinez FJ, Nielsen CM, Lias AM, King LDW, Edwards NJ, Greenwood NM, Kingham L, Poulton ID, Khozoee B, Goh C, Hodgson SH, Mac Lochlainn DJ, Salkeld J, Guillotte-Blisnick M, Huon C, Mohring F, Reimer JM, Chauhan VS, Mukherjee P, Biswas S, Taylor IJ, Lawrie AM, Cho J-S, Nugent FL, Long CA, Moon RW, Miura K, Silk SE, Chitnis CE, Minassian AM, Draper SJ. 2023. Vaccination with Plasmodium vivax Duffy-binding protein inhibits parasite growth during controlled human malaria infection. Science Translational Medicine 15:eadf1782.

20. Singh K, Mukherjee P, Shakri AR, Singh A, Pandey G, Bakshi M, Uppal G, Jena R, Rawat A, Kumar P, Bhardwaj R, Yazdani SS, Hans D, Mehta S, Srinivasan A, Anil K, Madhusudhan RL, Patel J, Singh A, Rao R, Gangireddy S, Patil R, Kaviraj S, Singh S, Carter D, Reed S, Kaslow DC, Birkett A, Chauhan VS, Chitnis CE. 2018. Malaria vaccine candidate based on Duffy-binding protein elicits strain transcending functional antibodies in a Phase I trial. npj Vaccines 3:48.

21. Payne RO, Silk SE, Elias SC, Milne KH, Rawlinson TA, Llewellyn D, Shakri AR, Jin J, Labbé GM, Edwards NJ, Poulton ID, Roberts R, Farid R, Jørgensen T, Alanine DG, de Cassan SC, Higgins MK, Otto TD, McCarthy JS, de Jongh WA, Nicosia A, Moyle S, Hill AV, Berrie E, Chitnis CE, Lawrie AM, Draper SJ. 2017. Human vaccination against Plasmodium vivax Duffy-binding protein induces strain-transcending antibodies. JCI insight. 2 (12):93683. doi:10.1172/jci.insight.93683.

22. Longley RJ, Sattabongkot J, Mueller I. 2016. Insights into the naturally acquired immune response to Plasmodium vivax malaria. Parasitology 143:154–170.

23. Rogers KJ, Vijay R, Butler NS. 2021. Anti-malarial humoral immunity: the long and short of it. Microbes Infect 23:104807.

24. Langhorne J, Ndungu FM, Sponaas AM, Marsh K. 2008. Immunity to malaria: more questions than answers. Nat Immunol 9:725–32.

25. Longley RJ, White MT, Takashima E, Brewster J, Morita M, Harbers M, Obadia T, Robinson LJ, Matsuura F, Liu ZSJ, Li-Wai-Suen CSN, Tham W-H, Healer J, Huon C, Chitnis CE, Nguitragool W, Monteiro W, Proietti C, Doolan DL, Siqueira AM, Ding XC, Gonzalez IJ, Kazura J, Lacerda M, Sattabongkot J, Tsuboi T, Mueller I. 2020. Development and validation of serological markers for detecting recent Plasmodium vivax infection. Nat Med 26:741–749.

26. Yman V, Tuju J, White MT, Kamuyu G, Mwai K, Kibinge N, Asghar M, Sundling C, Sondén K, Murungi L, Kiboi D, Kimathi R, Chege T, Chepsat E, Kiyuka P, Nyamako L, Osier FHA, Färnert A. 2022. Distinct kinetics of antibodies to 111 Plasmodium falciparum proteins identifies markers of recent malaria exposure. Nat Commun 13:331.

27. Lu F, Xu J, Liu Y, Ren Z, Chen J, Gong W, Yin Y, Li Y, Qian L, He X, Han X, Lin Z, Lu J, Zhang W, Liu J, Menard D, Han E-T, Cao J. 2024. Plasmodium vivax serological exposure markers: PvMSP1-42-induced humoral and memory B-cell response generates long-lived antibodies. PLOS Pathogens 20:e1012334.

28. Le Roch KG, Johnson JR, Florens L, Zhou Y, Santrosyan A, Grainger M, Yan SF, Williamson KC, Holder AA, Carucci DJ, Yates JR, 3rd, Winzeler EA. 2004. Global analysis of transcript and protein levels across the Plasmodium falciparum life cycle. Genome Res 14:2308–18.

29. Foth BJ, Zhang N, Chaal BK, Sze SK, Preiser PR, Bozdech Z. 2011. Quantitative Time-course Profiling of Parasite and Host Cell Proteins in the Human Malaria Parasite *Plasmodium falciparum*\*. Molecular & Cellular Proteomics 10.

30. Hazzard B, Sá JM, Ellis AC, Pascini TV, Amin S, Wellems TE, Serre D. 2022. Long read single cell RNA sequencing reveals the isoform diversity of Plasmodium vivax transcripts. bioRxiv doi:10.1101/2022.07.14.500005.

31. Arévalo-Herrera M, Lopez-Perez M, Dotsey E, Jain A, Rubiano K, Felgner PL, Davies DH, Herrera S. 2016. Antibody Profiling in Naïve and Semi-immune Individuals Experimentally Challenged with Plasmodium vivax Sporozoites. PLOS Neglected Tropical Diseases 10:e0004563.

32. Chuquiyauri R, Molina DM, Moss EL, Wang R, Gardner MJ, Brouwer KC, Torres S, Gilman RH, Llanos-Cuentas A, Neafsey DE, Felgner P, Liang X, Vinetz JM. 2015. Genome-Scale Protein Microarray Comparison of Human Antibody Responses in Plasmodium vivax Relapse and Reinfection. Am J Trop Med Hyg 93:801–809.

33. Doolan DL, Mu Y, Unal B, Sundaresh S, Hirst S, Valdez C, Randall A, Molina D, Liang X, Freilich DA, Oloo JA, Blair PL, Aguiar JC, Baldi P, Davies DH, Felgner PL. 2008. Profiling humoral immune responses to P. falciparum infection with protein microarrays. Proteomics 8:4680–4694.

34. Crompton PD, Kayala MA, Traore B, Kayentao K, Ongoiba A, Weiss GE, Molina DM, Burk CR, Waisberg M, Jasinskas A, Tan X, Doumbo S, Doumtabe D, Kone Y, Narum DL, Liang X, Doumbo OK, Miller LH, Doolan DL, Baldi P, Felgner PL, Pierce SK. 2010. A prospective analysis of the Ab response to *Plasmodium falciparum* before and after a malaria season by protein microarray. Proceedings of the National Academy of Sciences 107:6958–6963.

35. Dent AE, Nakajima R, Liang L, Baum E, Moormann AM, Sumba PO, Vulule J, Babineau D, Randall A, Davies DH, Felgner PL, Kazura JW. 2015. Plasmodium falciparum Protein Microarray Antibody Profiles Correlate With Protection From Symptomatic Malaria in Kenya. The Journal of Infectious Diseases 212:1429–1438.

36. Szymczak LC, Kuo H-Y, Mrksich M. 2018. Peptide arrays: Development and application. Anal Chem 90:266–282.

37. Buus S, Rockberg J, Forsström B, Nilsson P, Uhlen M, Schafer-Nielsen C. 2012. High-resolution Mapping of Linear Antibody Epitopes Using Ultrahigh-density Peptide Microarrays * . Molecular & Cellular Proteomics 11:1790–1800.

38. Kringelum JV, Nielsen M, Padkjær SB, Lund O. 2013. Structural analysis of B-cell epitopes in antibody:protein complexes. Mol Immunol 53:24–34.

39. Dahlbäck M, Rask TS, Andersen PH, Nielsen MA, Ndam NT, Resende M, Turner L, Deloron P, Hviid L, Lund O, Pedersen AG, Theander TG, Salanti A. 2006. Epitope mapping and topographic analysis of VAR2CSA DBL3X involved in P. falciparum placental sequestration. PLoS Pathog 2:e124.

40. Jaenisch T, Heiss K, Fischer N, Geiger C, Bischoff FR, Moldenhauer G, Rychlewski L, Sié A, Coulibaly B, Seeberger PH, Wyrwicz LS, Breitling F, Loeffler FF. 2019. High-density Peptide Arrays Help to Identify Linear Immunogenic B-cell Epitopes in Individuals Naturally Exposed to Malaria Infection. Mol Cell Proteomics 18:642–656.

41. Friedman-Klabanoff DJ, Travassos MA, Ifeonu OO, Agrawal S, Ouattara A, Pike A, Bailey JA, Adams M, Coulibaly D, Lyke KE, Laurens MB, Takala-Harrison S, Kouriba B, Kone AK, Doumbo OK, Patel JJ, Thera MA, Felgner PL, Tan JC, Plowe CV, Berry AA. 2021. Epitope-Specific Antibody Responses to a Plasmodium falciparum Subunit Vaccine Target in a Malaria-Endemic Population. J Infect Dis 223:1943–1947.

42. Bailey JA, Berry AA, Travassos MA, Ouattara A, Boudova S, Dotsey EY, Pike A, Jacob CG, Adams M, Tan JC, Bannen RM, Patel JJ, Pablo J, Nakajima R, Jasinskas A, Dutta S, Takala-Harrison S, Lyke KE, Laurens MB, Niangaly A, Coulibaly D, Kouriba B, Doumbo OK, Thera MA, Felgner PL, Plowe CV. 2020. Microarray analyses reveal strain-specific antibody responses to Plasmodium falciparum apical membrane antigen 1 variants following natural infection and vaccination. Sci Rep 10:3952.

43. Friedman-Klabanoff DJ, Jensen TL, Lyke KE, Laurens MB, Silva JC, Stucke EM, Ouattara A, Ifeonu OO, Hodges T, Moser KA, Gelber CE, Goll JB, Hoffman SL, Patel JJ, Pinapati RS, Tan JC, Deye GA, Takala-Harrison S, Travassos MA, Berry AA. 2025. Controlled human malaria infection with NF54 and 7G8 strains elicit differential antibody responses to Plasmodium falciparum peptides. Front Immunol 16:1641280.

44. Mitran CJ, Higa LM, Good MF, Yanow SK. 2020. Generation of a Peptide Vaccine Candidate against Falciparum Placental Malaria Based on a Discontinuous Epitope. Vaccines (Basel) 8.

45. Popovici J, Friedrich LR, Kim S, Bin S, Run V, Lek D, Cannon MV, Menard D, Serre D. 2018. Genomic Analyses Reveal the Common Occurrence and Complexity of Plasmodium vivax Relapses in Cambodia. MBio 9.

46. Grimée M, Tacoli C, Sandfort M, Obadia T, Taylor AR, Vantaux A, Robinson LJ, Lek D, Longley RJ, Mueller I, Popovici J, White MT, Witkowski B. 2024. Using serological diagnostics to characterize remaining high-incidence pockets of malaria in forest-fringe Cambodia. Malar J 23:49.

47. Auburn S, Böhme U, Steinbiss S, Trimarsanto H, Hostetler J, Sanders M, Gao Q, Nosten F, Newbold CI, Berriman M, Price RN, Otto TD. 2016. A new Plasmodium vivax reference sequence with improved assembly of the subtelomeres reveals an abundance of pir genes. Wellcome Open Res 1:4.

48. Xue B, Dunbrack RL, Williams RW, Dunker AK, Uversky VN. 2010. PONDR-FIT: a meta-predictor of intrinsically disordered amino acids. Biochim Biophys Acta 1804:996–1010.

49. Kim A, Popovici J, Menard D, Serre D. 2019. Plasmodium vivax transcriptomes reveal stage-specific chloroquine response and differential regulation of male and female gametocytes. Nat Commun 10:371.

50. Kim D, Paggi JM, Park C, Bennett C, Salzberg SL. 2019. Graph-based genome alignment and genotyping with HISAT2 and HISAT-genotype. Nat Biotechnol 37:907–915.

51. Singer M, Kanatani S, Castillo SG, Frischknecht F, Sinnis P. 2024. The Plasmodium circumsporozoite protein. Trends Parasitol 40:1124–1134.

52. Galaway F, Drought LG, Fala M, Cross N, Kemp AC, Rayner JC, Wright GJ. 2017. P113 is a merozoite surface protein that binds the N terminus of Plasmodium falciparum RH5. Nat Commun 8:14333.

53. Niaré K, Chege T, Rosenkranz M, Mwai K, Saßmannshausen Z, Odera D, Nyamako L, Tuju J, Alfred T, Waitumbi JN, Ogutu B, Sirima SB, Awandare G, Kouriba B, Rayner JC, Osier FHA. 2023. Characterization of a novel Plasmodium falciparum merozoite surface antigen and potential vaccine target. Front Immunol 14:1156806.

54. Garza R, Marchioni JM, Honeycutt JD, Hurlburt NK, Torres C, Garcia A, Loranc E, Yemington E, Towers D, Ssewanyana I, Pancera M, Lavinder JJ, Jagannathan P, Greenhouse B, Bol S, Bunnik EM. 2025. The N-terminal region of malaria vaccine candidate *Plasmodium falciparum* asparagine-rich merozoite antigen is immunodominant and targeted by polyreactive antibodies doi:10.64898/2025.12.11.693633. openRxiv.

55. Calvo-Calle JM, Mitchell R, Altszuler R, Othoro C, Nardin E. 2021. Identification of a neutralizing epitope within minor repeat region of Plasmodium falciparum CS protein. npj Vaccines 6:10.

56. Jelínková L, Flores-Garcia Y, Shapiro S, Roberts BT, Petrovsky N, Zavala F, Chackerian B. 2022. A vaccine targeting the L9 epitope of the malaria circumsporozoite protein confers protection from blood-stage infection in a mouse challenge model. npj Vaccines 7:34.

57. Flores-Garcia Y, Wang LT, Park M, Asady B, Idris AH, Kisalu NK, Muñoz C, Pereira LS, Francica JR, Seder RA, Zavala F. 2021. The P. falciparum CSP repeat region contains three distinct epitopes required for protection by antibodies in vivo. PLoS Pathog 17:e1010042.

58. Ferreira MU, Nunes MdS, Wunderlich G. 2004. Antigenic Diversity and Immune Evasion by Malaria Parasites. Clinical and Vaccine Immunology 11:987–995.

59. Davies HM, Nofal SD, McLaughlin EJ, Osborne AR. 2017. Repetitive sequences in malaria parasite proteins. FEMS Microbiol Rev 41:923–940.

60. Belachew EB. 2018. Immune Response and Evasion Mechanisms of Plasmodium falciparum Parasites. J Immunol Res 2018:6529681.

61. Muralidharan V, Goldberg DE. 2013. Asparagine Repeats in Plasmodium falciparum Proteins: Good for Nothing? PLOS Pathogens 9:e1003488.

62. van der Lee R, Buljan M, Lang B, Weatheritt RJ, Daughdrill GW, Dunker AK, Fuxreiter M, Gough J, Gsponer J, Jones DT, Kim PM, Kriwacki RW, Oldfield CJ, Pappu RV, Tompa P, Uversky VN, Wright PE, Babu MM. 2014. Classification of Intrinsically Disordered Regions and Proteins. Chemical Reviews 114:6589–6631.

63. Verra F, Hughes AL. 1999. Biased amino acid composition in repeat regions of Plasmodium antigens. Mol Biol Evol 16:627–33.

64. Murugan R, Scally SW, Costa G, Mustafa G, Thai E, Decker T, Bosch A, Prieto K, Levashina EA, Julien JP, Wardemann H. 2020. Evolution of protective human antibodies against Plasmodium falciparum circumsporozoite protein repeat motifs. Nat Med 26:1135–1145.

65. Beeson JG, Drew DR, Boyle MJ, Feng G, Fowkes FJI, Richards JS. 2016. Merozoite surface proteins in red blood cell invasion, immunity and vaccines against malaria. FEMS Microbiol Rev 40:343–372.

66. Malpede BM, Tolia NH. 2014. Malaria adhesins: structure and function. Cell Microbiol 16:621–31.

67. Wang Q, Zhao Z, Zhang X, Li X, Zhu M, Li P, Yang Z, Wang Y, Yan G, Shang H, Cao Y, Fan Q, Cui L. 2016. Naturally Acquired Antibody Responses to Plasmodium vivax and Plasmodium falciparum Merozoite Surface Protein 1 (MSP1) C-Terminal 19 kDa Domains in an Area of Unstable Malaria Transmission in Southeast Asia. PLoS One 11:e0151900.

68. Nogueira PA, Alves FP, Fernandez-Becerra C, Pein O, Santos NR, Pereira da Silva LH, Camargo EP, del Portillo HA. 2006. A reduced risk of infection with Plasmodium vivax and clinical protection against malaria are associated with antibodies against the N terminus but not the C terminus of merozoite surface protein 1. Infect Immun 74:2726–33.

69. Patel PN, Dickey TH, Hopp CS, Diouf A, Tang WK, Long CA, Miura K, Crompton PD, Tolia NH. 2022. Neutralizing and interfering human antibodies define the structural and mechanistic basis for antigenic diversion. Nat Commun 13:5888.

70. Soares IS, da Cunha MG, Silva MN, Souza JM, Del Portillo HA, Rodrigues MM. 1999. Longevity of naturally acquired antibody responses to the N- and C-terminal regions of Plasmodium vivax merozoite surface protein 1. Am J Trop Med Hyg 60:357–63.

71. Zuo S, Lu J, Sun Y, Song J, Han S, Feng X, Han ET, Cheng Y. 2024. The Plasmodium vivax MSP1P-19 is involved in binding of reticulocytes through interactions with the membrane proteins band3 and CD71. J Biol Chem 300:107285.

72. Pizarro JC, Normand BV-L, Chesne-Seck M-L, Collins CR, Withers-Martinez C, Hackett F, Blackman MJ, Faber BW, Remarque EJ, Kocken CHM, Thomas AW, Bentley GA. 2005. Crystal Structure of the Malaria Vaccine Candidate Apical Membrane Antigen 1. Science 308:408–411.

73. Treeck M, Zacherl S, Herrmann S, Cabrera A, Kono M, Struck NS, Engelberg K, Haase S, Frischknecht F, Miura K, Spielmann T, Gilberger TW. 2009. Functional analysis of the leading malaria vaccine candidate AMA-1 reveals an essential role for the cytoplasmic domain in the invasion process. PLoS Pathog 5:e1000322.

74. Richard D, MacRaild CA, Riglar DT, Chan J-A, Foley M, Baum J, Ralph SA, Norton RS, Cowman AF. 2010. Interaction between Plasmodium falciparum apical membrane antigen 1 and the rhoptry neck protein complex defines a key step in the erythrocyte invasion process of malaria parasites. J Biol Chem 285:14815–14822.

75. Shen B, Sibley LD. 2012. The moving junction, a key portal to host cell invasion by apicomplexan parasites. Curr Opin Microbiol 15:449–455.

76. Lamarque M, Besteiro S, Papoin J, Roques M, Vulliez-Le Normand B, Morlon-Guyot J, Dubremetz J-F, Fauquenoy S, Tomavo S, Faber BW, Kocken CH, Thomas AW, Boulanger MJ, Bentley GA, Lebrun M. 2011. The RON2-AMA1 interaction is a critical step in moving junction-dependent invasion by apicomplexan parasites. PLoS Pathog 7:e1001276.

77. Maskus DJ, Królik M, Bethke S, Spiegel H, Kapelski S, Seidel M, Addai-Mensah O, Reimann A, Klockenbring T, Barth S, Fischer R, Fendel R. 2016. Characterization of a novel inhibitory human monoclonal antibody directed against Plasmodium falciparum Apical Membrane Antigen 1. Sci Rep 6:39462.

78. Patel PN, Diouf A, Dickey TH, Tang WK, Hopp CS, Traore B, Long CA, Miura K, Crompton PD, Tolia NH. 2025. A strain-transcending anti-AMA1 human monoclonal antibody neutralizes malaria parasites independent of direct RON2L receptor blockade. Cell Rep Med 6:101985.

79. Winnicki AC, Dietrich MH, Yeoh LM, Carias LL, Roobsoong W, Drago CL, Malachin AN, Redinger KR, Feufack-Donfack LB, Baldor L, Jung NC, McLaine OS, Skomorovska-Prokvolit Y, Orban A, Opi DH, Kirtley P, Nielson K, Aleshnick M, Zanghi G, Rezakhani N, Vaughan AM, Wilder BK, Sattabongkot J, Tham W-H, Popovici J, Beeson JG, Bosch J, King CL. 2024. Potent AMA1-specific human monoclonal antibody against Plasmodium vivax Pre-erythrocytic and Blood Stages. Nature Communications 15:10556.

80. Patel PN, Dickey TH, Diouf A, Salinas ND, McAleese H, Ouahes T, Long CA, Miura K, Lambert LE, Tolia NH. 2023. Structure-based design of a strain transcending AMA1-RON2L malaria vaccine. Nat Commun 14:5345.

81. Olivieri A, Collins CR, Hackett F, Withers-Martinez C, Marshall J, Flynn HR, Skehel JM, Blackman MJ. 2011. Juxtamembrane Shedding of Plasmodium falciparum AMA1 Is Sequence Independent and Essential, and Helps Evade Invasion-Inhibitory Antibodies. PLOS Pathogens 7:e1002448.

82. Srinivasan P, Beatty WL, Diouf A, Herrera R, Ambroggio X, Moch JK, Tyler JS, Narum DL, Pierce SK, Boothroyd JC, Haynes JD, Miller LH. 2011. Binding of Plasmodium merozoite proteins RON2 and AMA1 triggers commitment to invasion. Proceedings of the National Academy of Sciences 108:13275–13280.

83. Zeeshan M, Tyagi RK, Tyagi K, Alam MS, Sharma YD. 2015. Host-parasite interaction: selective Pv-fam-a family proteins of Plasmodium vivax bind to a restricted number of human erythrocyte receptors. J Infect Dis 211:1111–1120.

84. Alam MS, Choudhary V, Zeeshan M, Tyagi RK, Rathore S, Sharma YD. 2015. Interaction of Plasmodium vivax tryptophan-rich antigen PvTRAg38 with Band 3 on human erythrocyte surface facilitates parasite growth. J Biol Chem 290:20257–20272.

85. Tyagi K, Hossain ME, Thakur V, Aggarwal P, Malhotra P, Mohmmed A, Sharma YD. 2016. Plasmodium vivax tryptophan rich antigen PvTRAg36.6 interacts with PvETRAMP and PvTRAg56.6 interacts with PvMSP7 during erythrocytic stages of the parasite. PLoS One 11:e0151065.

86. Wang B, Lu F, Cheng Y, Chen J-H, Jeon H-Y, Ha K-S, Cao J, Nyunt MH, Han J-H, Lee S-K, Kyaw MP, Sattabongkot J, Takashima E, Tsuboi T, Han E-T. 2015. Immunoprofiling of the Tryptophan-Rich Antigen Family in Plasmodium vivax. Infection and Immunity 83:3083–3095.

87. Kundu P, Naskar D, McKie SJ, Dass S, Kanjee U, Introini V, Ferreira MU, Cicuta P, Duraisingh M, Deane JE, Rayner JC. 2023. The structure of a Plasmodium vivax Tryptophan Rich Antigen domain suggests a lipid binding function for a pan-Plasmodium multi-gene family. Nat Commun 14:5703.

88. Alam MS, Rathore S, Tyagi RK, Sharma YD. 2016. Host-parasite interaction: multiple sites in the Plasmodium vivax tryptophan-rich antigen PvTRAg38 interact with the erythrocyte receptor band 3. FEBS Lett 590:232–241.

89. Hester J, Chan ER, Menard D, Mercereau-Puijalon O, Barnwell J, Zimmerman PA, Serre D. 2013. De novo assembly of a field isolate genome reveals novel Plasmodium vivax erythrocyte invasion genes. PLoS Negl Trop Dis 7:e2569.

90. Ntumngia FB, Thomson-Luque R, Torres LdM, Gunalan K, Carvalho LH, Adams JH. 2016. A Novel Erythrocyte Binding Protein of Plasmodium vivax Suggests an Alternate Invasion Pathway into Duffy-Positive Reticulocytes. MBio 7.

91. Lee S-K, Crosnier C, Valenzuela-Leon PC, Dizon BLP, Atkinson JP, Mu J, Wright GJ, Calvo E, Gunalan K, Miller LH. 2024. Complement receptor 1 is the human erythrocyte receptor for Plasmodium vivax erythrocyte binding protein. Proc Natl Acad Sci U S A 121:e2316304121.

92. Chootong P, Ntumngia FB, VanBuskirk KM, Xainli J, Cole-Tobian JL, Campbell CO, Fraser TS, King CL, Adams JH. 2010. Mapping Epitopes of the *Plasmodium vivax* Duffy Binding Protein with Naturally Acquired Inhibitory Antibodies. Infection and Immunity 78:1089–1095.

93. He W-Q, Shakri AR, Bhardwaj R, França CT, Stanisic DI, Healer J, Kiniboro B, Robinson LJ, Guillotte-Blisnick M, Huon C, Siba P, Cowman A, King CL, Tham W-H, Chitnis CE, Mueller I. 2019. Antibody responses to Plasmodium vivax Duffy binding and Erythrocyte binding proteins predict risk of infection and are associated with protection from clinical Malaria. PLoS Negl Trop Dis 13:e0006987.

94. King CL, Michon P, Shakri AR, Marcotty A, Stanisic D, Zimmerman PA, Cole-Tobian JL, Mueller I, Chitnis CE. 2008. Naturally acquired Duffy-binding protein-specific binding inhibitory antibodies confer protection from blood-stage *Plasmodium vivax* infection. Proceedings of the National Academy of Sciences 105:8363–8368.

95. Urusova D, Carias L, Huang Y, Nicolete VC, Popovici J, Roesch C, Salinas ND, Dechavanne S, Witkowski B, Ferreira MU, Adams JH, Gross ML, King CL, Tolia NH. 2019. Structural basis for neutralization of Plasmodium vivax by naturally acquired human antibodies that target DBP. Nat Microbiol 4:1486–1496.

96. Ntumngia FB, Schloegel J, Barnes SJ, McHenry AM, Singh S, King CL, Adams JH. 2012. Conserved and Variant Epitopes of Plasmodium vivax Duffy Binding Protein as Targets of Inhibitory Monoclonal Antibodies. Infection and Immunity 80:1203–1208.

97. Chen E, Salinas ND, Ntumngia FB, Adams JH, Tolia NH. 2015. Structural analysis of the synthetic Duffy Binding Protein (DBP) antigen DEKnull relevant for Plasmodium vivax malaria vaccine design. PLoS Negl Trop Dis 9:e0003644.

98. Chan L-J, Dietrich MH, Nguitragool W, Tham W-H. 2020. Plasmodium vivax Reticulocyte Binding Proteins for invasion into reticulocytes. Cell Microbiol 22:e13110.

99. Nagaoka H, Kanoi BN, Jinoka K, Morita M, Arumugam TU, Palacpac NMQ, Egwang TG, Horii T, Tsuboi T, Takashima E. 2019. The N-terminal region of Plasmodium falciparum MSP10 is a target of protective antibodies in malaria and is important for PfGAMA/PfMSP10 interaction. Front Immunol 10:2669.

100. Kehrer J, Kuss C, Andres-Pons A, Reustle A, Dahan N, Devos D, Kudryashev M, Beck M, Mair GR, Frischknecht F. 2018. Nuclear pore complex components in the malaria parasite Plasmodium berghei. Sci Rep 8:11249.

101. Dahan-Pasternak N, Nasereddin A, Kolevzon N, Pe’er M, Wong W, Shinder V, Turnbull L, Whitchurch CB, Elbaum M, Gilberger TW, Yavin E, Baum J, Dzikowski R. 2013. PfSec13 is an unusual chromatin-associated nucleoporin of Plasmodium falciparum that is essential for parasite proliferation in human erythrocytes. J Cell Sci 126:3055–3069.

102. Stanisic DI, Good MF. 2023. Malaria vaccines: Progress to date. BioDrugs 37:737–756.

103. Teixeira LH, Tararam CA, Lasaro MO, Camacho AGA, Ersching J, Leal MT, Herrera S, Bruna-Romero O, Soares IS, Nussenzweig RS, Ertl HCJ, Nussenzweig V, Rodrigues MM. 2014. Immunogenicity of a prime-boost vaccine containing the circumsporozoite proteins of Plasmodium vivax in rodents. Infect Immun 82:793–807.

104. Parobek CM, Bailey JA, Hathaway NJ, Socheat D, Rogers WO, Juliano JJ. 2014. Differing patterns of selection and geospatial genetic diversity within two leading Plasmodium vivax candidate vaccine antigens. PLoS Negl Trop Dis 8:e2796.

105. White MT, Karl S, Battle KE, Hay SI, Mueller I, Ghani AC. 2014. Modelling the contribution of the hypnozoite reservoir to Plasmodium vivax transmission. eLife 3:e04692.

106. Combe A, Moreira C, Ackerman S, Thiberge S, Templeton TJ, Ménard R. 2009. TREP, a novel protein necessary for gliding motility of the malaria sporozoite. Int J Parasitol 39:489–496.

107. Bauza K, Malinauskas T, Pfander C, Anar B, Jones EY, Billker O, Hill AVS, Reyes-Sandoval A. 2014. Efficacy of a Plasmodium vivax Malaria Vaccine Using ChAd63 and Modified Vaccinia Ankara Expressing Thrombospondin-Related Anonymous Protein as Assessed with Transgenic Plasmodium berghei Parasites. Infection and Immunity 82:1277–1286.

108. Cabral-Miranda G, Heath MD, Mohsen MO, Gomes AC, Engeroff P, Flaxman A, Leoratti FMS, El-Turabi A, Reyes-Sandoval A, Skinner MA, Kramer MF, Bachmann MF. 2017. Virus-Like Particle (VLP) Plus Microcrystalline Tyrosine (MCT) Adjuvants Enhance Vaccine Efficacy Improving T and B Cell Immunogenicity and Protection against Plasmodium berghei/vivax. Vaccines 5:10.

109. Kariu T, Yuda M, Yano K, Chinzei Y. 2002. MAEBL Is Essential for Malarial Sporozoite Infection of the Mosquito Salivary Gland. Journal of Experimental Medicine 195:1317–1323.

110. Sá M, Costa DM, Teixeira AR, Pérez-Cabezas B, Formaglio P, Golba S, Sefiane-Djemaoune H, Amino R, Tavares J. 2022. MAEBL Contributes to Plasmodium Sporozoite Adhesiveness. International journal of molecular sciences. 23 (10):5711. doi:10.3390/ijms23105711.

111. Cunningham D, Lawton J, Jarra W, Preiser P, Langhorne J. 2010. The pir multigene family of Plasmodium: antigenic variation and beyond. Mol Biochem Parasitol 170:65–73.

